# Backbone-mediated weakening of pairwise interactions enables percolation in peptide-based mimics of protein condensates

**DOI:** 10.1101/2024.07.08.602299

**Authors:** Xiangze Zeng, Rohit V. Pappu

**Affiliations:** Department of Physics, Hong Kong Baptist University, Kowloon Tong, Hong Kong and Teaching and Research Division, School of Chinese Medicine, Hong Kong Baptist University, Kowloon Tong, Hong Kong; Department of Biomedical Engineering and Center for Biomolecular Condensates, The James McKelvey School of Engineering, Washington University in St. Louis, St. Louis, MO 63130, USA

## Abstract

Biomolecular condensates formed by intrinsically disordered proteins (IDPs) condensates are semidilute solutions. These can be approximated as solutions of blob-sized segments, which can be as small as peptide-sized motifs. We leveraged the blob picture to quantify differences between inter-residue interactions in model compound and peptide-based mimics of dense versus dilute phases. The all-atom molecular dynamics simulations use the polarizable AMOEBA forcefield. In model compound solutions, the interactions between aromatic residues are stronger than interactions between cationic and aromatic residues. This holds in dilute and dense phases. Cooperativity within dense phases enhances pairwise interactions leading to finite-sized nanoscale clusters. The results for peptide-based condensates paint a different picture. Backbone amides add valence to the associating molecules. While this maintains or enhances pairwise inter-residue interactions in dilute phases, it weakens pair interactions in dense phases. Weakening of pair interactions enables fluidization characterized by short-range order and long-range disorder. The higher valence afforded by the peptide backbone becomes a generator of system-spanning networks. As a result, dense phases of peptides are best described as percolated network fluids. Overall, our results show how peptide backbones enhance pairwise interactions in dilute phases whole weakening these interactions in dense phases to enable percolation within dense phases.

## INTRODUCTION

Phase separation of proteins and nucleic acids leads to the formation of biomolecular condensates^1^. In a two-component system comprising a protein and solvent, phase separation yields two coexisting phases namely, a dilute phase that is deficient in protein and a coexisting dense phase that is enriched in protein ^2^. Intrinsically disordered regions (IDRs) of proteins that feature specific sequence characteristics ^3^ can be drivers of protein phase separation ^4^. Of relevance to condensates such as stress granules ^5^ and P-bodies ^6^ are IDRs known as prion-like low complexity domains (PLCDs) ^7^. The sequence determinants of driving forces for phase separation of PLCDs have been the topic of several studies^4, 8, 9, 10, 11, 12, 13, 14, 15, 16, 17, 18, 19, 20, 21, 22, 23, 24, 25^.

Recently, Bremer et al. quantified the temperature-dependent driving forces for phase separation of ∼40 different sequence variants of A1-LCD, which is the PLCD from hnRNP-A1 ^25^. At a given temperature, each A1-LCD variant is defined by a saturation concentration *c*_sat_, which is the threshold concentration for phase separation. Lower values of *c*_sat_ imply stronger driving forces for phase separation and vice versa. The temperature-specific values of *c*_sat_ were found to be governed by: (i) the number and types of aromatic residues (Phe versus Tyr), which are uniformly distributed along the linear sequence ^8, 26^; (ii) the context-dependent contributions of arginine residues; (iii) the types of polar residues, which in PLCDs appear to function as spacers that are interspersed between aromatic and Arg stickers; (iv) the destabilizing effects of lysine residues that interfere with π-π interactions; and (v) the contributions of net charge per residue. Away from the critical temperature, changes to sequence features can lead to changes in temperature-specific values of *c*_sat_ that can span 4-5 orders of magnitude ^25^. In contrast, while the material properties of dense phases formed by variants of A1-LCD, specifically the measured and computed viscosities, show a clear inverse correlation with *c*_sat_ ^27^, the concentrations within dense phases vary by less than a factor of 2. These findings suggest that the material properties of dense phases and the values of *c*_sat_ are driven primarily by the free energy of transfer of PLCDs from dilute to dense phases ^22^. The results also imply that interactions in dilute versus dense phases must be very different from one another ^28^.

Dense phases of PLCDs and other systems are equivalent to semidilute solutions of flexible linear polymers (**Figure 1**) ^10, 25^. This designation is based on the concentrations within dense phases which are above the overlap concentration and well below the values expected for a polymer melt. Furthermore, several simulations and recent measurements have shown that the sizes (radii of gyration and mean end-to-end distances) of IDRs are expanded within condensates when compared to coexisting dilute phases ^8, 9, 10, 29, 30, 31, 32, 33, 34, 35, 36^. The excluded volumes appear to be positive within condensates and this suggests that condensate interiors are akin to good solvents or at least better than theta solvents for which the excluded volume is zero ^9, 10^. For systems such as A1-LCD and a series of sequence variants Farag et al. showed that the radii of gyration within dense phases change minimally with temperature ^10^. The measured concentrations within dense phases also change minimally, by less than a factor 2, across the entire temperature range that approaches the critical point ^10, 25^. Hence, it can be argued that condensate interiors are athermal and or good solvents for systems such as PLCDs ^37^.

**Figure 1.**
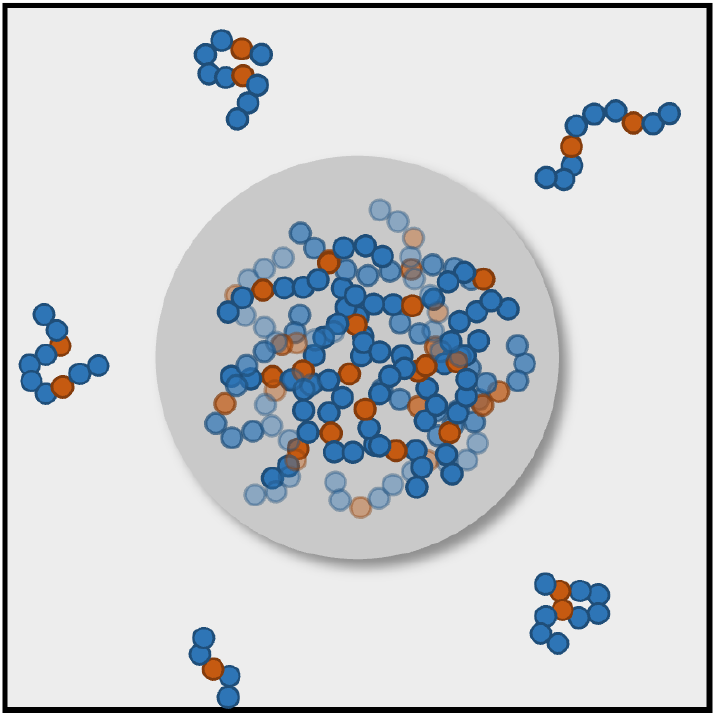
Dense phases of protein-based condensates are semidilute solutions of flexible linear polymers. ^10^. Here, the bulk volume fraction of polymers is at or larger than the overlap volume fraction. The overlap volume fraction is defined as the ratio of the occupied volume of the polymer chain v_mono_*N* to the pervaded volume *V*, where *N* is the number of monomer in one polymer chain and v_mono_ is the occupied volume of one chemical monomer. *V* can be roughly estimated as *R*^3^, where *R* is the size of the chain. The blue spheres denote spacers, and the orange spheres denote stickers.

In semidilute solutions of flexible polymers in good or athermal solvents, the polymer mass concentration *c* is at or above the overlap mass concentration *c*\* ^28^. This allows the use of the concept of concentration blobs that was introduced by de Gennes ^38^. Each polymer is viewed as a chain of blobs. The size ξ_c_ of a blob scales as g_c_^3/5^, where g_c_ is the number of residues within a blob. Beyond the blob, chain sizes in semidilute solutions scale as (n/g_c_)^1/2^, where n is the number of residues in the chain. Correlations among blobs decay as (*c*/*c*\*)^-3/4^. Importantly, values of g_c_ do not depend on n ^37, 39^. Instead, they depend purely on the excluded volume, v_ex_ ^39^. In an athermal solvent, v_ex_ is at its upper limit, and g_c_ = 1. Given published results ^10, 25^, v_ex_ appears to be near the upper limit, and hence g_c_ is ≈ 1. Accordingly, as a zeroth-order approximation, one can approximate dense phases as fluids of peptide-sized segments in aqueous solvents ^40^.

Here, we leverage the blob picture of de Gennes and report results from molecular dynamics simulations for capped amino acids (1 < g_c_ < 2) and for capped tripeptides (3 < g_c_ < 4). Peptides encompass backbone amides and sidechains. To dissect the contributions of backbones and sidechains, we compare the interactions of systems comprising sidechains alone to systems comprising backbones and sidechains. The blob picture suggests that our simulations that use capped amino acids or capped tripeptides will have a direct bearing on understanding how interactions are affected within condensate interiors. In the all-atom, molecular dynamics simulations, we investigated the concentration dependence of the strengths of effective pairwise interactions among functional groups that are known to be important as drivers of phase separation of PLCDs ^8, 9, 10, 25, 27^. Specifically, our simulations quantify the strengths of interactions among aromatic and cationic residues in mimics of dilute versus dense phases. Volume fractions of sidechain model compounds between 0.002 and 0.003 were used to mimic the dilute phase whereas volume fractions between 0.08 and 0.09 mimic the dense phase.

For the simulations, we used the polarizable forcefield, AMOEBA ^41, 42, 43^, which has been parameterized using high-level quantum calculations, validated extensively, and widely used to: (a) investigate protein conformational changes; (b) examine ion solvation ^44^; (c) model the structural and thermodynamic properties of water ^45, 46^; (d) compute hydration free energies of peptides and small molecules ^47^; (e) obtain absolute and relative alchemical free energies ^48^; (f) simulate protein conformational equilibria ^49, 50, 51^; (g) perform free energy calculations of protein-ligand binding ^49, 52, 53^; (h) simulate DNA aptamers with ligands ^54^; (i) calculate electric fields in liquid solutions; and (j) model RNA hybridization ^52^. For our work, we used the AMOEBA parameters for protein atoms, water molecules, and solution ions ^41^.

We used a polarizable forcefield to allow for the formal possibility that the interiors of dense phases might be different from those of dilute phases. Structural studies based on super resolution microscopy have shown that dense phases feature spatial inhomogeneities defined by nanoscale hubs that coexist with dispersed molecules that create networks of meshes of different length sizes ^55^. These findings are concordant with inferences from lattice-based simulations ^10, 55^. Inhomogeneities in condensate interiors are manifest as nanoscale clustering of macromolecules and this is enabled by physical crosslinking ^10, 56^. The internal dynamics of clusters and dispersed molecules reflect the making and breaking of crosslinks on disparate timescales ^55, 56^. Recent studies have also pointed to condensates having distinct chemical and electrochemical environments ^12^. Furthermore, recent THz measurements revealed the presence of two types of water populations within condensates ^57^. These populations are governed by the types of species that are being hydrated. Structural and solvation inhomogeneities within condensates have been used in direct computations and explanations of condensate viscoelasticity ^27, 56, 58, 59, 60, 61, 62, 63, 64^. In light of these rapidly emerging findings and given that condensates are likely to be inhomogeneously organized and heterogeneously crowded, we used a polarizable forcefield to allow for the ability to capture first-order electrostatic responses that cannot be captured using non-polarizable forcefields. Our results show that even though the thermodynamic inferences do not qualitatively change vis-à-vis non-polarizable forcefields, there is a material impact on the self-diffusion coefficient of water molecules. These results are consistent with findings from THz measurements and other reports ^57, 65^.

The design of our simulations is as follows: For different pairs of amino acid sidechains, we performed two sets of simulations. In the first set, we used model compound mimics of specific amino acid side chains (**Figure 2A**) to compute potentials of mean force (PMFs) and quantify the free energies of association / dissociation between functional groups of sidechains. To assess dilute phase interactions among distinct motifs, we computed PMFs between pairs of model compounds in a cubic box of volume ∼125 nm^3^. Each model compound represents a specific amino acid sidechain. To quantify interactions among sidechains in dense phases, we computed PMFs from simulations of mixtures of model compounds as a function of concentration.

**Figure 2.**
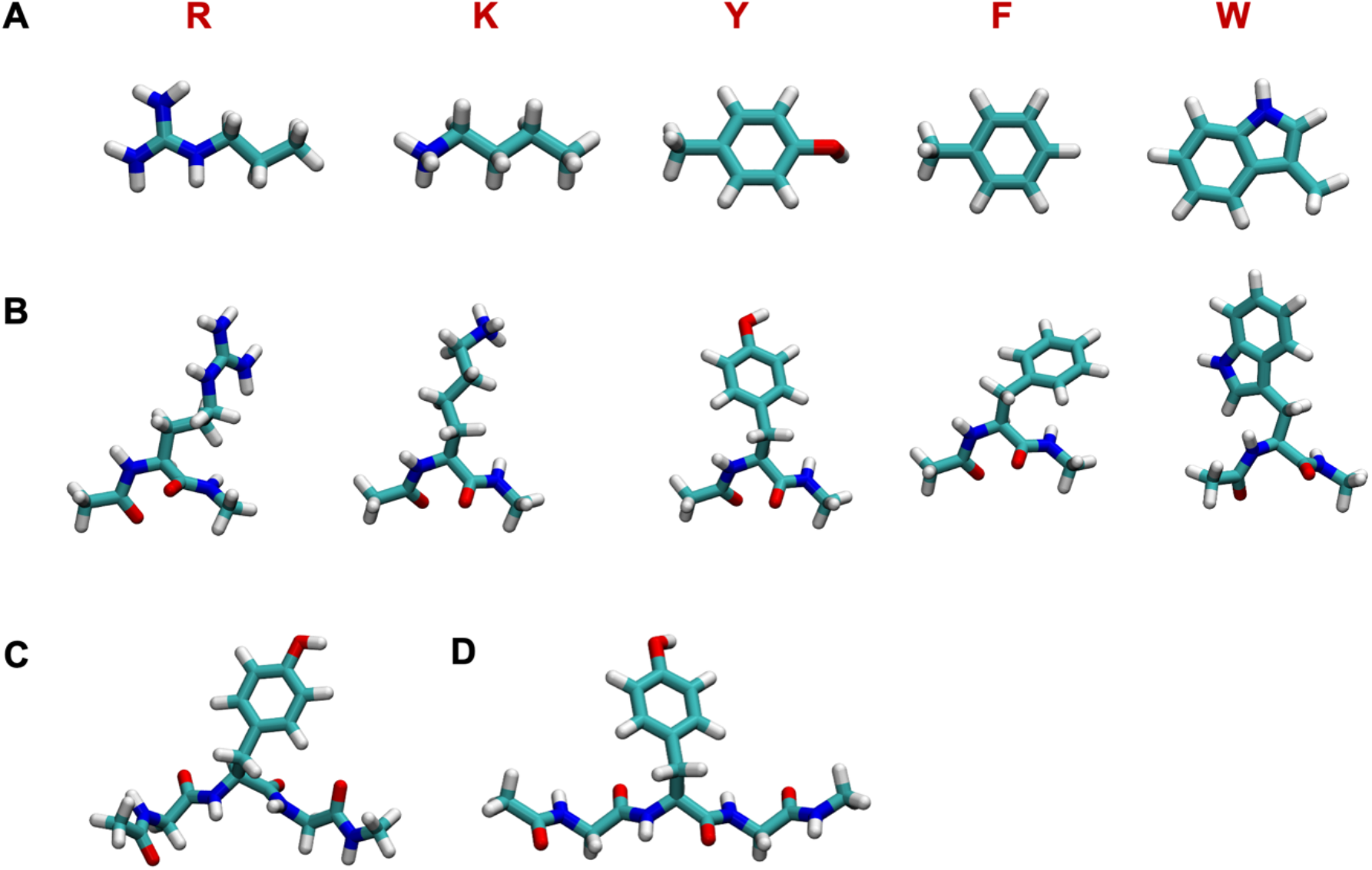
Model compounds and capped amino acids used to compare interactions between pairs of polypeptide units in mimics of dilute versus dense phases. (A) Five model compounds *n*-propylguanidine (R′), 1-butylamine (K′), *p*-Cresol (Y′), toluene (F′), and 3-methylindole (W′) that mimic the side chains of Arg, Lys, Tyr, Phe, and Trp, respectively. (B) Drawings of capped amino acids, N-acetyl-Xaa-N′-methylamide for Xaa ≡ Arg, Lys, Tyr, Phe and Trp, respectively. Each of these systems feature two peptide units and one pair of backbone ϕ and ψ angles. We also performed simulations for a fully flexible tripeptide (C) N-acetyl-Gly-Tyr-Gly-N′-methylamide GYG and (D) the tripeptide N-acetyl- Gly-Tyr-Gly -N′-methylamide restrained to be in an extended conformation.

In a second set of simulations, we computed inter-sidechain PMFs using dilute and dense phase simulations of capped amino acids (**Figure 2B**). Here, a capped amino acid Xaa refers to the molecule N-acetyl-Xaa-N′-methylamide, where Xaa is the amino acid residue of interest. Each capped amino acid features two peptide units, and a pair of unrestrained backbone ϕ and ψ angles.

Tripeptides were also used to test sequence context effects and conformational contributions (**Figure 2C, 2D**). Comparative assessments of the simulations of model compound solutions and those of capped amino acids and tripeptides help us query the concentration-dependent effects of backbone peptide units and local sequence contexts on the strengths of sidechain interactions.

## RESULTS

### PMFs in dilute phase follow well-defined hierarchies

We computed dilute phase PMFs for pairs of model compounds that mimic the following interactions: Phe-Phe, Tyr-Tyr, Trp-Trp, Arg-Phe, Arg-Tyr, Arg-Trp, Lys-Phe, and Lys-Tyr. The model compounds that mimic sidechains of Arg, Lys, Tyr, Phe and Trp are *n*-propylguanidine (R′), 1-butylamine (K′), *p*-Cresol (Y′), toluene (F′), and 3-methylindole (W′), respectively (**Figure 2A**). These model compounds were chosen because they are the dominant drivers or destabilizers of PLCD phase separation ^9, 10, 25^.

For the dilute phase, we computed PMFs for different pairs of model compounds using umbrella sampling. Volume fractions of sidechain model compounds between 0.002 and 0.003 were used to mimic the dilute phase. Each pair was solvated in a cubic box of water and solution ions, ∼5 nm to a side, with periodic boundary conditions. The reaction coordinate chosen for umbrella sampling was the distance between specific atoms on functional groups of each model compound. For R′ and K′, the chosen atoms were the carbon atom in the guanidium group of Arg and the nitrogen atom in the amine of Lys. For Y′, F′, and W′, we used the geometric center of the aromatic ring to define the reaction coordinate.

The PMFs for each of the six model compound pairs are shown in **Figures 3A-3C**. The PMF for Ý-Ý shows two minima at ∼3.6 Å and ∼5.0 Å, respectively (**Figure 3A**). These minima correspond to two different interaction modes, namely the stacked structure and T-shape structure (**Figure S1**). The locations of these minima are consistent with results reported for benzene dimers using high-level quantum calculations ^66^. The PMFs for F′-F′ and W′-W′ have one minimum each, with the minimum at ∼5.0 Å for F′-F′ corresponding to the T-shaped structure, and the minimum at ∼3.6 Å for W′-W′ corresponding to the stacked structure (**Figure 3A**). The dominant minima in PMFs for R′-Y′, R′-F′, and R′-W′ appear at a separation of ∼3.6 Å and this corresponds to stacked structures formed by the planar guanidinium group and the rings of the aromatic groups (**Figure S2**). These structures are consistent with reports of interactions between Arg and aromatic residues being of cation-π-π flavor ^67^. We also computed PMFs for K′-Y′ and K′-F′. These interactions are essentially negligible when compared to those of R′-Y′ and R′-F′ (**Figure 3C**).

**Figure 3.**
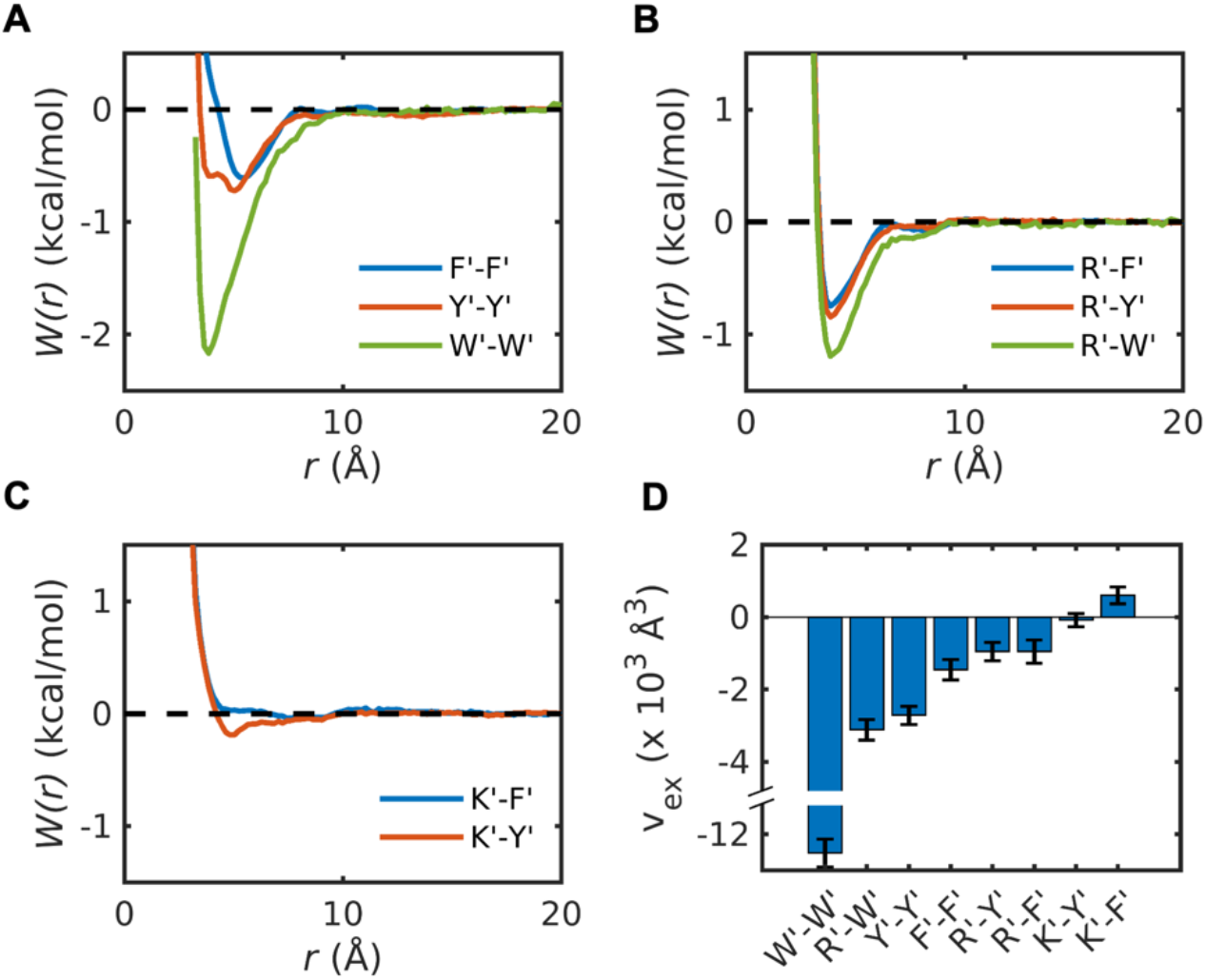
PMFs computed for pairs of model compound mimics of sidechains. PMFs, *W*(*r*), for interactions between π systems Phe, Tyr, and Trp, (B) Arg-π systems, and (C) Lys-π systems. The model compounds that mimic sidechains of Arg, Lys, Tyr, Phe and Trp are represented by R′, K′, Y′, F′, and W′, respectively. (D) Each of the pairwise PMFs can be used to compute pair-specific excluded volumes (v_ex_), as described in the text.

To enable a single-parameter comparison of the strengths and types of pairwise interactions, we computed excluded volumes (v_ex_) by integrating the Mayer *f*-function ^68^. For this, we computed 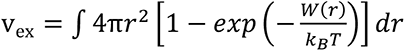. Here, (𝑟) is the PMF for pairwise interactions computed from the simulations, and 4π*r*^2^*dr* is the volume element. The value of v_ex_, which quantifies the volume excluded for interactions with the solvent ^68, 69^, is the two-body interaction coefficient between a pair of moieties in solvent and it is directly proportional to the second virial coefficient ^2, 68^. A negative value implies that the interactions are on average attractive, whereas positive values imply net repulsions. The magnitude of v_ex_ quantifies the strength of the attraction or repulsion.

The values of v_ex_ computed using the PMFs are shown in **Figure 3D**. The strengths of attractive attractions, quantified by negative values of v_ex_, follow the hierarchy W′-W′ > R′-W′ > Y′-Y′ > F′-F′ > R′-Y′ > R’-F′. The v_ex_ values for K′-Y′ and K′-F′ are near zero (K′-Y′) or positive, albeit small (K′-F′), implying that these interactions are negligible or slightly repulsive. These results align with experiments demonstrating that R′-Y′ interactions are stronger than R′-F′ interactions and K′-Y′ interactions ^4, 8, 25, 70, 71, 72^. Note that the direct interactions between water molecules and model compounds are weaker than the solvent-mediated pair interactions between model compounds (**Figure S3**).

Next, we computed dilute phase PMFs between pairs of capped amino acids along reaction coordinates that are identical to those for model compounds (**Figure 4**). The reaction coordinates were identical to those used for model compounds. This enables a one-to-one correspondence between the two sets of simulations.

**Figure 4.**
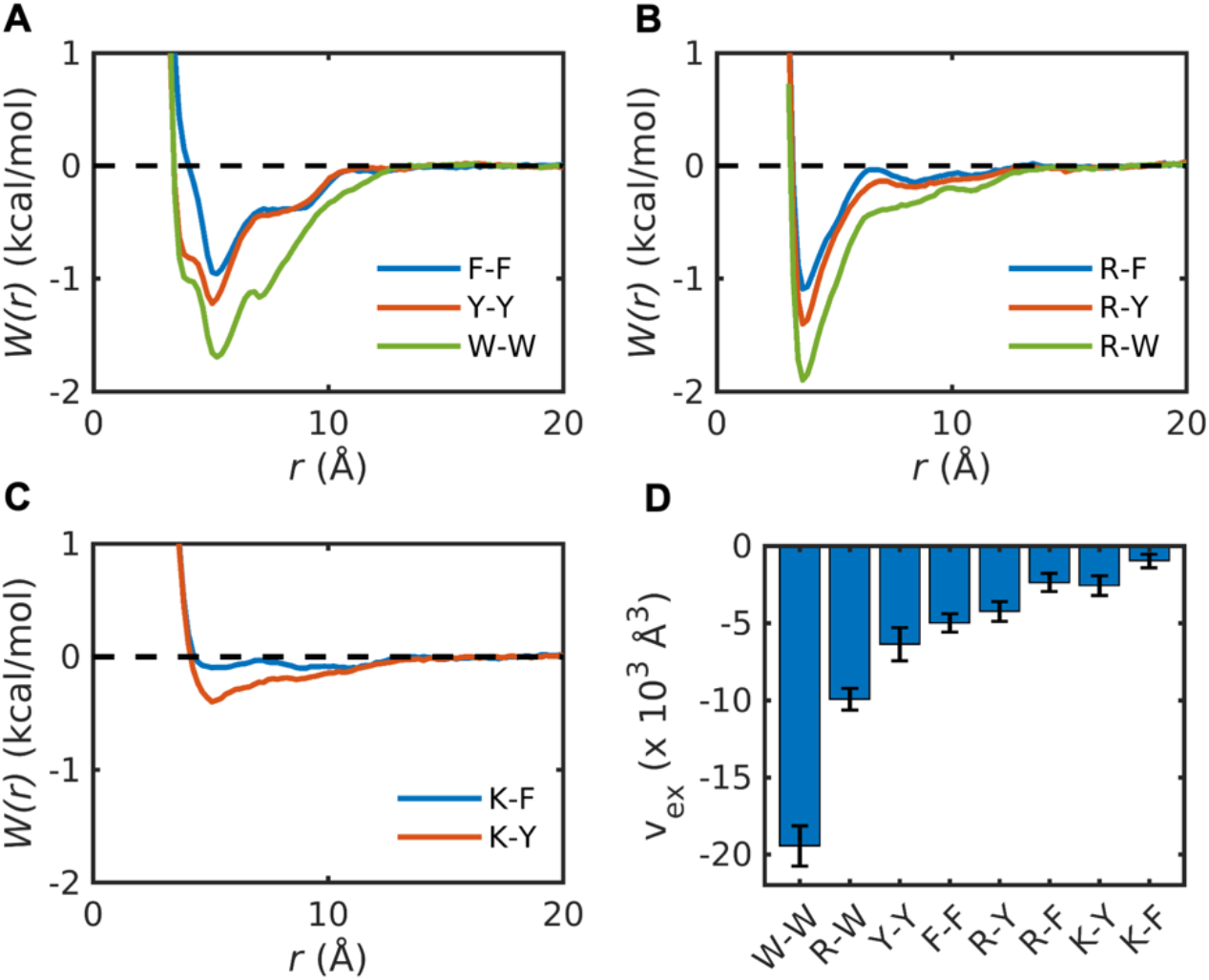
Interactions between sidechains are enhanced by the presence of backbone peptide units in the dilute phase. PMFs, *W*(*r*), for (A) π-π, (B) Arg-π, and (C) Lys-π interactions between pairs of capped amino acids. (D) The pairwise PMFs in panels (A) – (C) were used to compute pair-specific excluded volumes (v_ex_).

Although the locations of minima remain the same as those for model compounds, the well depths increase for all sidechain pairs. Additionally, auxiliary minima appear at separations of ∼8.5 Å for all pairs. These point to backbone-mediated, long-range contacts between functional groups of sidechains that are absent for model compounds. Although backbones enhance the overall interaction strengths between sidechains, the overall hierarchy remains the same as can be seen by comparing the excluded volumes in **Figure 4D** to those in **Figure 3D**. To quantify the influence of backbones on the interactions between side chains, we calculated Δ(𝑟) = 𝑊_AA_(𝑟) − 𝑊_mc_(r). This refers to the difference between PMFs for capped amino *W*_AA_(*r*) those for model compounds 𝑊_mc_(r) (**Figure S4**). These comparisons show that backbones help enhance attractions between sidechains in dilute phases.

We quantified the entropic and enthalpic contributions to the PMFs. The potential energy was used as the proxy for enthalpy, and its averaged value was plotted as a function of the distance *r* (**Figure S11**). Despite significant fluctuations, the results show that entropy is favorable at short range within 10 Å for W-W and F-F interactions. These findings highlight the importance of hydrophobic hydration to the association of aromatic rings and align with the findings reported by Grimme ^73^ as well as Martinez and Iverson ^74^. Both studies propose that the π-π stacking among small aromatic rings, that is observed in high-resolution structures, should be used as a geometrical description rather than as a marker of special interactions compared to those between saturated hydrocarbon rings. They also showed that the interaction strengths of π-π stacking among small aromatic rings are almost identical to those between saturated hydrocarbon rings of similar size. For other pairs, enthalpy dominates the interactions at distances *r* < ∼4Å.

### Model compounds form nanoscale clusters within dense phases

Next, we quantified the effective pairwise interaction strengths among sidechains in dense phase mimics made up of model compounds. In these simulations, the volume fractions of sidechains are more than an order of magnitude larger than in the dilute phase, ranging from 0.08 to 0.09. These values correspond to measured volume fractions of sidechains that place PLCD-like systems in two-phase regimes^25^. The corresponding concentrations for each of these systems are shown in **Table S1**. We performed seven separate simulations for systems comprising different numbers of: R′ and F′, R′ and Y′, R′ and W′, respectively. For each pair, we performed two or three sets of simulations. In 1:1 mixtures, the simulations comprised 40 R and 40 F′ / Y′ / W′ molecules (**Figure 5A**). In 2:1 mixtures the simulations comprised 53 R′ and 27 F′ / Y′ molecules (**Figure 5B**). Finally, in 1:2 mixtures, the simulations comprised 27 R′ and 53 F′ / Y′ molecules (**Figure 5C**).

**Figure 5.**
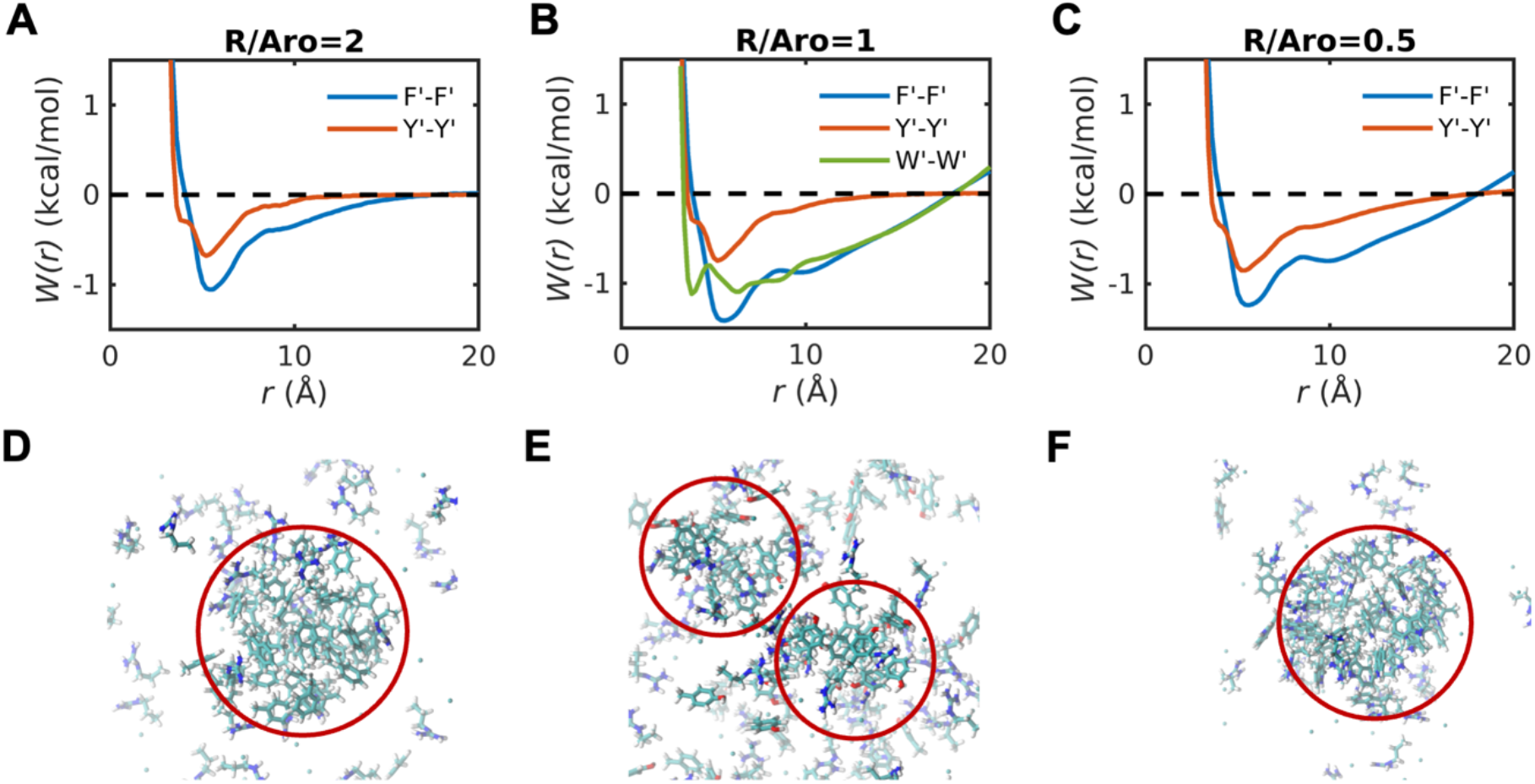
**Strengths of interactions between aromatic groups are enhanced in dense phases of model compounds.** (A) PMFs, *W*(*r*), for F′-F′ and Y′-Y′ in 1:1 mixtures of R′ and F′, and. Each mixture includes 53 R′ and 27 aromatic (either F′ or Y′) sidechains. (B) *W*(*r*) for F′-F′, Y′-Y′ and W′-W′ in 1:1 mixtures of Arg model compound with each aromatic model compound. Each mixture includes 40 Arg and 40 aromatic (either F′, Y′ or W′) model compounds. (C) *W*(*r*) for F′-F′ and Y′-Y′ in 1:2 mixtures of Arg model compound with each aromatic model compound. Each mixture includes 27 Arg and 53 aromatic (either F′ or Y′) model compounds. Snapshots of nanoscale clusters that are formed in 1:1 mixtures of R with (D) F′, (E) Y′, and (F) W′.

When compared to the dilute phase PMFs (**Figure 3A**), the PMFs for the dense phase mimics (**Figure 5A-5C**) show the following trends: The energetics for Y-Y interactions change the least, with the only substantive change being the increased preference for T-shaped structures. When the R:Y ratio is 2:1 and 1:1, the PMFs converge to zero for separations between 10 Å and 15 Å. This indicates an absence of long-range ordering in R-Y mixtures (**Figure 5A-5C**). However, when the R:Y ratio is 1:2, then we observe long-range ordering that goes beyond 20 Å.

The PMFs for F′-F′ change both qualitatively and quantitatively. When compared to the dilute phase PMFs (**Figure 3A**), the T-shaped structures have higher stability in dense phases. Additionally, we observe long-range, non-random organization for 1:1 and 2:1 ratios of R′:Y′ (**Figures 5A** and **5C**). This is indicative of clustering on the nanometer scale. Increasing the R′:F′ ratio weakens the preference for T-shaped structures and long-range ordering (**Figure 5B**). Decreasing the R′:F′ ratio has the opposite effect (**Figure 5C**).

In R′:W′ mixtures, the PMF for W′-W′ interactions show an equivalent preference for stacked and T-shaped structures (**Figure 5A**), which is different from the PMFs in the dilute phase (**Figure 3A**). This weakens the preference for stacked structures when compared to the dilute phase. In addition to the equivalent preferences of stacked and T-shaped structures, the 1:1 R′-W′ mixtures show the long-range ordering we observe for R-F mixtures. Snapshots derived from simulations of 1:1 mixtures show the formation of nanoscale clusters of aromatic moieties in R′-

F′ (**Figure 5D**) and R′-W′ (**Figure 5F**) mixtures, and weaker clustering in R′-Y′ mixtures (**Figure 5E**).

The weakened clustering of aromatic moieties in R′-Y′ mixtures can be rationalized by comparisons of the R′-Y′, R′-F′, and R′-W′ PMFs. The minimum, which corresponds to stacked structures of the guanido group and aromatic moieties, is most pronounced in R′-Y′ mixtures and is weaker in R′-F′ and R′-W′ mixtures (**Figure 6A**). The extent to which R′-F′ interactions are weakened depends on the R′:F′ ratio, becoming more pronounced as this ratio decreases (**Figure 6B**). In contrast, the R′-Y′ interactions, although destabilized when compared to the dilute phase, are insensitive to changes in the R′:Y′ ratio in dense phase mimics (**Figure 6C**). These results suggest a correlation between stronger R′-Y′ interactions and weaker clustering of Tyr mimics, whereas the converse is true for R′-F′ and R′-W′ mixtures. In contrast to R′-F′ and R′-Y′ mixtures, the interactions in K′-F′ and K′-Y′ mixtures are such that interactions are either negligible (K′-F′, **Figure 6B**) or weakly attractive (K′-Y′, **Figure 6C**). These interactions in dense phases, which do not depend on concentration or the K′:F′ / K′:Y′ ratios, resemble what is observed in dilute phases (**Figure 3C**).

**Figure 6.**
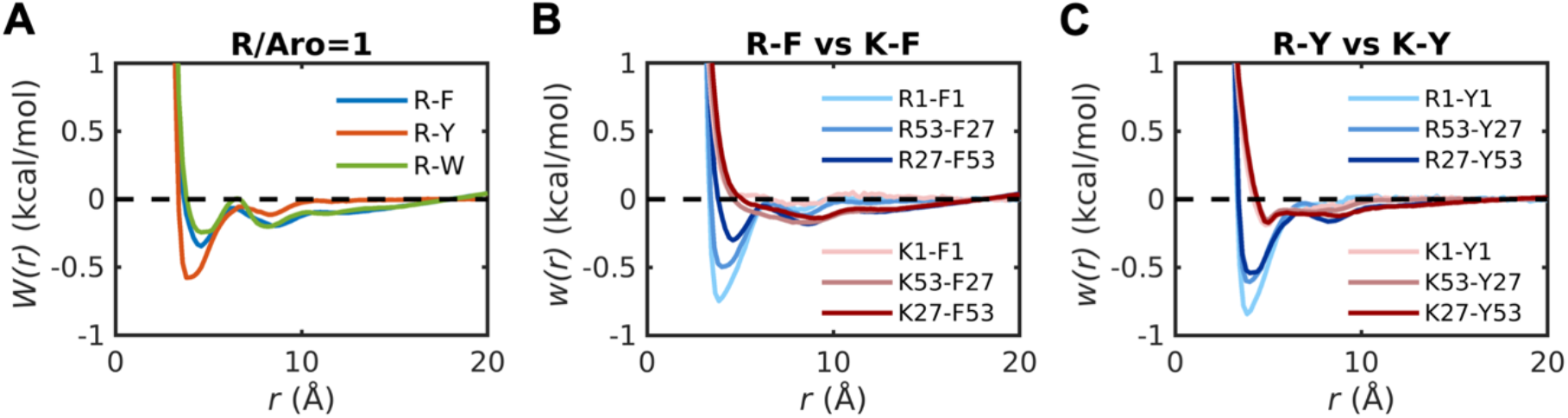
Comparative strengths of cation-π interactions in dense phases with different ratios of. **R**′**, K**′ **and aromatic (Aro) groups.** (A) PMFs, *W*(*r*), for R′-F′, R-Y, and R′-W′ in 1:1 mixtures of R′ with each aromatic model compound. Each mixture includes 40 R′ molecules and 40 molecules of one of F′, Y′ or W′. (B) Comparison of R′-F′ and K′-F′ in different dense phases. R1-F1 and K1-F1 denote *W*(*r*) for R′-F′ and K′-F′ in the dilute phase calculated from umbrella sampling. The other legends denote *W*(*r*) in mixtures containing different numbers of model compounds mimicking Arg and aromatic sidechains. In the legend, R53-F27 refers to PMFs extracted from simulations of dense phases with 53 R′ and 27 F′ molecules. Conversely, R27-F53 refers to PMFs extracted from simulations of dense phases with 27 R′ and 53 F′ molecules. A similar convention is applied for PMFs computed in dense phases with different mixtures of K′ and F′ molecules. (C) Comparison of R′-Y′ and K′-Y′ in different dilute and dense phases. Conventions for the legends are the same as those in panel (B).

We also explored the influence of total model compound concentration while maintaining a 1:1 ratio of *n*-propylguanidine to aromatic moieties (**Figure S5**). The effects we observed for Y′- Y′, F′-F′, R′-Y′, and R′-F′ interactions (**Figure 5** and **Figure 6**) become more pronounced with increasing concentration (**Figure S5**). Nanoscale clusters of Tyr moieties, which are absent at lower concentrations, are manifest in mixtures with 60 R′ and 60 Y′ molecules. This implies that the formation of nanoscale aromatic clusters is a generic feature of dense solutions with aromatic molecules. However, the specificity of cluster formation, which depends on the solvent- and mixture-mediated strengths of associations among aromatic moieties, will be manifest in the concentration dependence of cluster formation. This point is made by the observation that nanoscale clusters form in 1:1 mixtures with 30 R′ and 30 F′ molecules.

### Pair interactions are weakened in dense phases of capped amino acids

Next, we performed simulations for mixtures of capped amino acids to compute pair PMFs for 1:1 mixtures of dense phase mimics of different combinations of capped Arg and aromatic residues (**Figure 7**). In all cases, we observed a weakening of pairwise associations and the extent of weakening increases within increasing concentrations. Furthermore, the pair PMFs for F-F, Y-F and W-W (**Figures 7A**, **7C**, and **7E**) do not show the long-range ordering that was observed for mixtures of model compounds (**Figure 5** and **Figure S5**).

**Figure 7.**
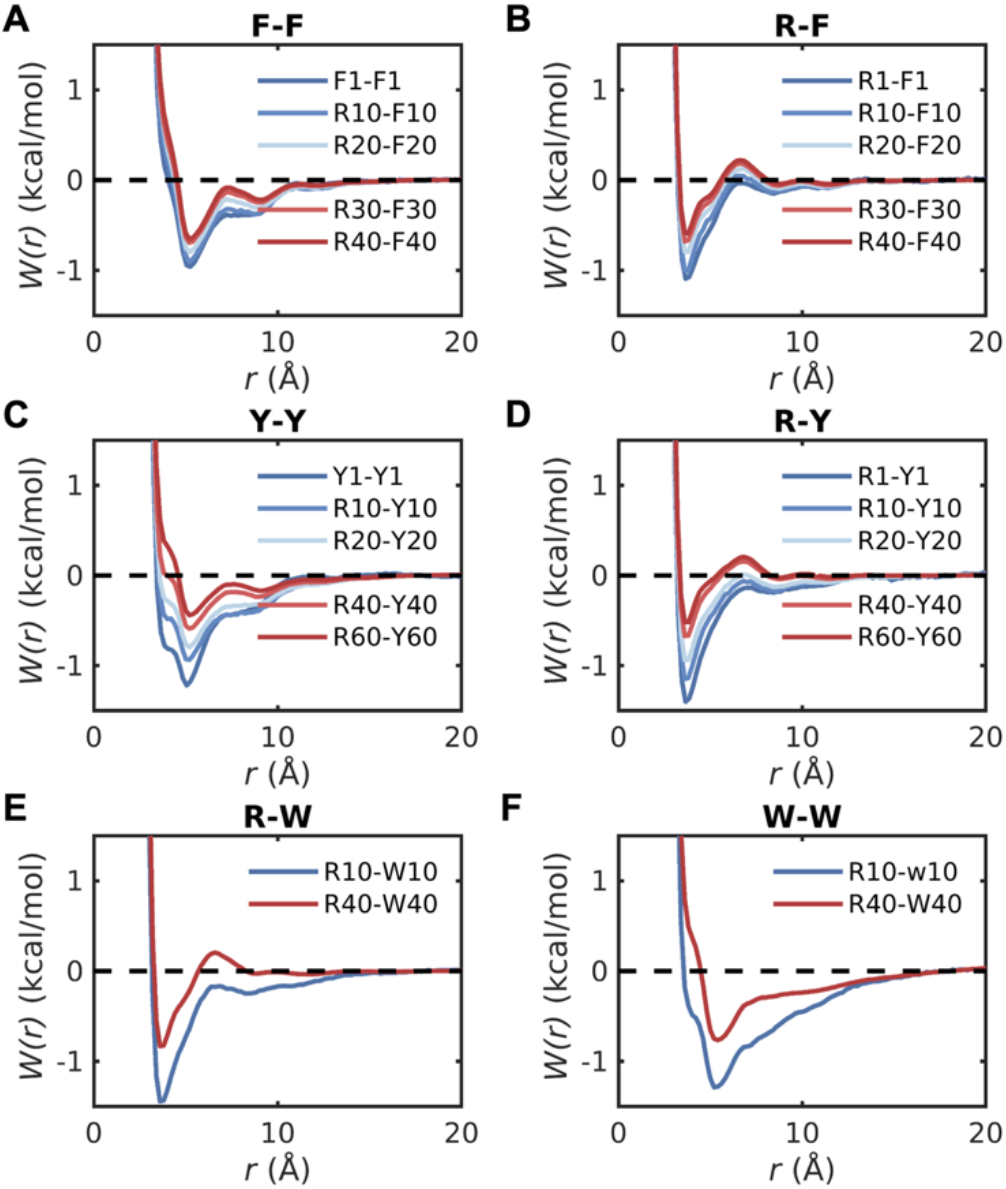
**Contributions from backbone peptide units weaken sidechain interactions within dense phases.** Comparisons of potentials of mean force *W*(*r*) in dilute versus dense phases for pairs of capped amino acids (A) F-F, (B) R-F, (C) Y-Y, (D) R-Y, (E) R-W, and (F) W-W. The number in the legends denotes the amount of model compounds indicated by the preceding letter in the mixture mimicking dense phases. F1-F1, Y1-Y1, R1-Y1, and R1- Y1 denote potential of mean force in dilute phase calculated from umbrella sampling. The interactions are weakened as increasing the concentration of amino acid stickers.

The information contained in the different pair PMFs is summarized by computing pair-specific values of v_ex_ for different concentrations of 1:1 mixtures of Arg and aromatic moieties in dense phase mimics based on model compounds (**Figures 8A-8B**) or capped amino acids (**Figures 8C-8D**). For the F-F, Y-Y, and W-W pairs, the magnitudes of v_ex_ increase in dense phase mimics that are based on the use of model compounds. However, for the same pairs of sidechains, the magnitudes of v_ex_ decrease with increasing concentrations of capped amino acids. Similar trends prevail when comparing the excluded volumes for pairwise associations of Arg and Tyr, Phe, or Trp (see **Figures 8B** versus **8D**). Overall, the pairwise associations are weakened, long-range attractions are lost, and nanoscale clusters are absent in dense phases of capped amino acids. The implication is that while the peptide unit enhances and maintains the hierarchy of pairwise associations observed in dilute phases, the interactions are fundamentally altered in dense phases. We asked if the weakening of pairwise associations is offset by a gain in sidechain-backbone or backbone-backbone pairwise associations. Analysis of the pair PMFs that quantify these pairwise associations show the same trends we observed for sidechain interactions (**Figure S6**). Furthermore, we found that the stoichiometry of Arg versus aromatic residues does not alter the PMFs for capped amino acids (**Figure S7**). This points to the robustness of the weakening of pairwise associations caused by the backbone-mediated interactions.

**Figure 8.**
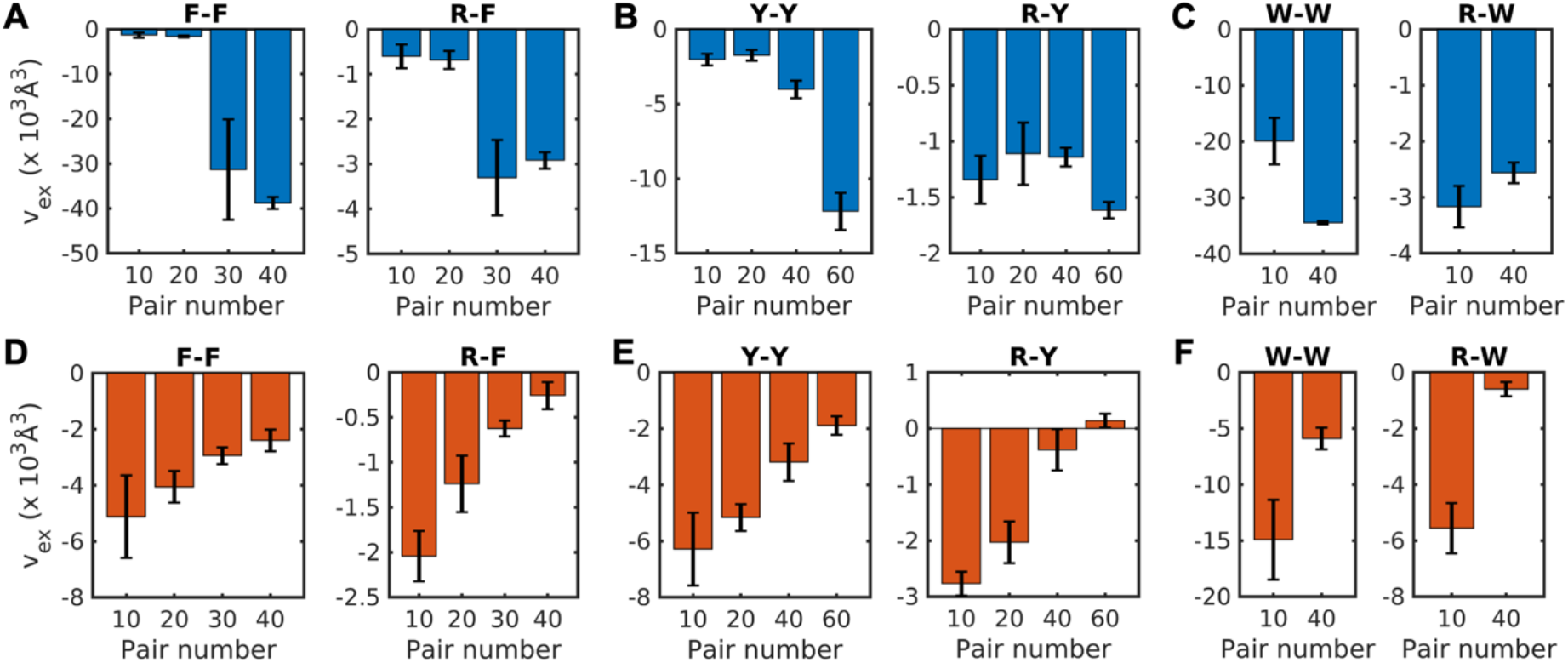
Pairwise interactions among sidechains are weakened in dense phases of amino acids. Excluded volumes (v_ex_) for pairwise interactions of model compounds (A)-(C) and capped amino acids (D)-(F). (A) F-F and R-F in 1:1 mixture of Arg and Phe model compounds, (B) Y-Y and R-Y in 1:1 mixture of Arg and Tyr model compounds, and (C) W-W and R-W in 1:1 mixture of Arg and Trp model compounds. (D) F-F and R-F in 1:1 mixture of Arg and Phe capped amino acids, (E) Y-Y and R-Y calculated in 1:1 mixture of Arg and Tyr capped amino acids, and (F) W-W and R-W in 1:1 mixture of Arg and Trp model compounds. X-axis represents the number of pairs of Arg and aromatic stickers in the mixture.

Next, we tested the effects of longer peptides on pairwise associations between Arg and aromatic residues, specifically Tyr. In PLCDs, aromatic residues are uniformly distributed along the linear sequence ^26^. Motifs such as GYG and GRG are common in PLCDs ^25^. Accordingly, we computed pair PMFs, with Arg-Tyr distances as the reaction coordinates, for two different combinations of capped GYG and GRG tripeptides. In one set of simulations, the backbone conformations were unrestrained. In the other set of simulations, the four pairs of backbone ϕ and ψ angles were restrained in extended β-strand-like conformations with values of ϕ = –140° and ψ = +135°.

Next, the PMFs for Y-Y associations computed in dense phase mimics with 40 GRG and 40 GYG peptides are compared to the PMFs for capped Arg and Tyr in dilute versus dense phases (**Figure 9A**). For the unrestrained pairs of tripeptides, the PMFs share the same profiles as those for capped amino acids, but the interactions are weakened in comparison. Restraining the backbone conformations, elicits a substantial entropic penalty, and this weakens pairwise associations between Tyr residues (**Figure 9A**). In all PMFs, the locations of stacked and T-shaped structures stay invariant, whereas the minimum corresponding to backbone-mediated associations, located at ∼ 8.5Å for capped amino acids and unrestrained tripeptide, shifts to ∼7.5Å for the restrained tripeptides. The PMFs that quantify R-Y associations are equivalent for capped amino acids and tripeptides **(Figure 9B**).

**Figure 9.**
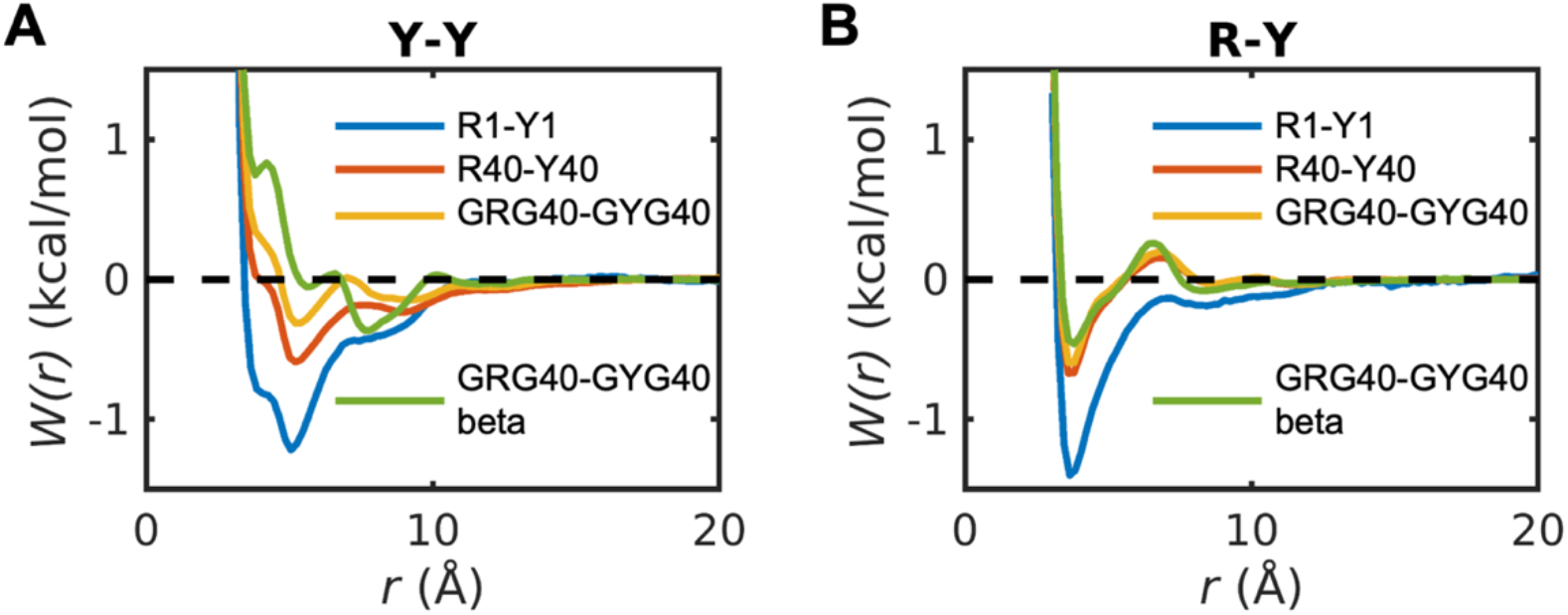
Influence of stoichiometry of cationic versus aromatic amino acids and backbone flexibility on pairwise interactions between capped amino acids. The potential of mean force for (A) Y-Y, and (B) R-Y in mixtures including capped Arg and Tyr. The number in the legends denotes the amount of model compounds indicated by the preceding letter in the mixture mimicking dense phases. Y1-Y1and R1-Y1 denote potential of mean force in dilute phase calculated from umbrella sampling. While the total concentration keeps the same, the stoichiometry of cationic vs. aromatic stickers does not affect the effective pairwise interactions between capped amino acids in the dense phase. The potential of mean force for (A) Y-Y, and (B) R-Y in mixtures including capped tripeptides GRG and GYG. The red curve denotes results when the backbone of tripeptide is constrained in the 𝛽 region where 𝜙 = −140°, 𝜓 = 135°.

The totality of the results presented to this point suggest that strengths of pairwise associations and long-range ordering due to strong pair interactions between aromatic residues are weakened by backbone-mediated effects. The loss of long-range ordering points to the fluidizing effects of backbone-mediated interactions. However, the pair associations present a conundrum. If they are the only associations to be considered, which is true in theories that rely exclusively on two-body interactions ^75^, then transferring peptides or proteins into dense phases should be thermodynamically unfavorable ^76^. The fact that phase separation is favored above system-specific threshold concentrations suggests that interactions beyond pairwise associations ^77^ must contribute to interactions within dense phases ^28, 78^. This point has been underscored recently to explain why compositionally distinct phases are likely to be determined by the contributions of three-body and higher-order interactions ^79^.

We investigated the contributions of higher-order interactions by quantifying the degree of clustering through connectivity ^80^, which is the basis for forming network fluids via directional interactions whereby molecules can be viewed as multivalent nodes connected to other molecules via edges on a graph ^40, 81^. This analysis was prompted by the fact that each peptide unit contributes at least one hydrogen bond donor and acceptor. Accordingly, the valence increases by four for capped amino acids and by right for capped tripeptides. Note that in mean field models, which are valid in semidilute solutions, any valence above three is sufficient to generate a percolated network ^2, 40, 68, 69, 81^. At or below a valence of three, small clusters can form, but they cannot grow into system-spanning networks ^69^.

### Dense phase mimics modeled using capped amino acids are percolated network fluids

We asked if the weakening of pairwise interactions and loss of long-range ordering in dense phases mimicked using capped amino acids are offset by a gain in higher-order interactions in the form of higher-order clusters and networks that span the entire system. To answer this question, we performed an analysis of the connectivity of molecules within dense phase mimics formed by capped amino acids ^40, 80, 81, 82^. The analysis was inspired by the recent work of Dar et al., ^81^. Two molecules were considered connected if any of the distances between specific atoms or groups is smaller than a prescribed cutoff. The pairwise distances used to compute connectivity were divided into three groups: (1) sidechain-sidechain, (2) sidechain-backbone, (3) and distances between backbone amide nitrogen and backbone carbonyl oxygen atoms. The inter-sidechain distances include those between aromatic rings, between guanidinium groups, and between the aromatic ring and guanidinium group. The sidechain-backbone distances include those between aromatic ring and terminal CH₃ groups, between aromatic ring and backbone carbonyl oxygen atoms, and between the guanidinium group and backbone carbonyl oxygen atoms. The geometric centers of carbon atoms in the aromatic ring, the nitrogen atom in the guanidinium group, and the carbon atom in the terminal CH_3_ group were used to calculate these distances. Following Dar et al., ^81^ the first minimum of the radial distribution function for a specific pair of atoms or groups was chosen as the distance cutoff to determine the connectivity. The values for these distance cutoffs are shown in **Table S2**. If molecule A is connected to molecules B and C, then molecule B and C are also connected via transitive closure ^83^. All connected molecules make up a cluster. We used this connectivity analysis to define the cluster size as the number of molecules within a cluster. We also defined the cluster dimension as the maximum distance between alpha carbon atoms within the cluster ^84^. For the cluster with only one molecule, we set the cluster dimension to be zero.

Results from the connectivity analysis are shown in **Figure 10**. The degree of connectivity, which quantifies the connectedness of a molecule within the dense phase, shows a monotonic increase with increasing concentration (**Figure 10A**). Percolation is a continuous transition ^40, 85^, whereby above a threshold the connectivity increases to span the entire system. The monotonic increase in the extent of clustering is also evidenced in the distributions of the number of clusters per molecule (**Figure 10B**). The distributions of cluster size show a bimodality at high concentrations, pointing to the incorporation of more than 75% of the molecules into the largest cluster. The largest cluster is system-spanning as evidenced by comparison to the maximum cluster dimension, which is equivalent to the dimension for the central simulation cell. The weakening of pairwise associations within dense phase mimics formed by capped amino acids is offset by the formation of higher-order connected clusters that become system-spanning at the highest concentrations. Unlike dense phases of capped amino acids, we do not observe system-spanning networks for dense phases made up of model compounds (**Figure S8**). Instead, these systems exhibit a lower degree of connectivity, smaller cluster dimensions, a higher number of clusters per molecule, and a clear failure to form system-spanning networks. Taken together with the absence of long-range ordering, the findings suggest that backbone-mediated interactions drive the formation of a percolated network within dense phases ^81^. This network has fluid-like characteristics defined by weakened pairwise associations, short-range order, and long-range disorder.

**Figure 10:**
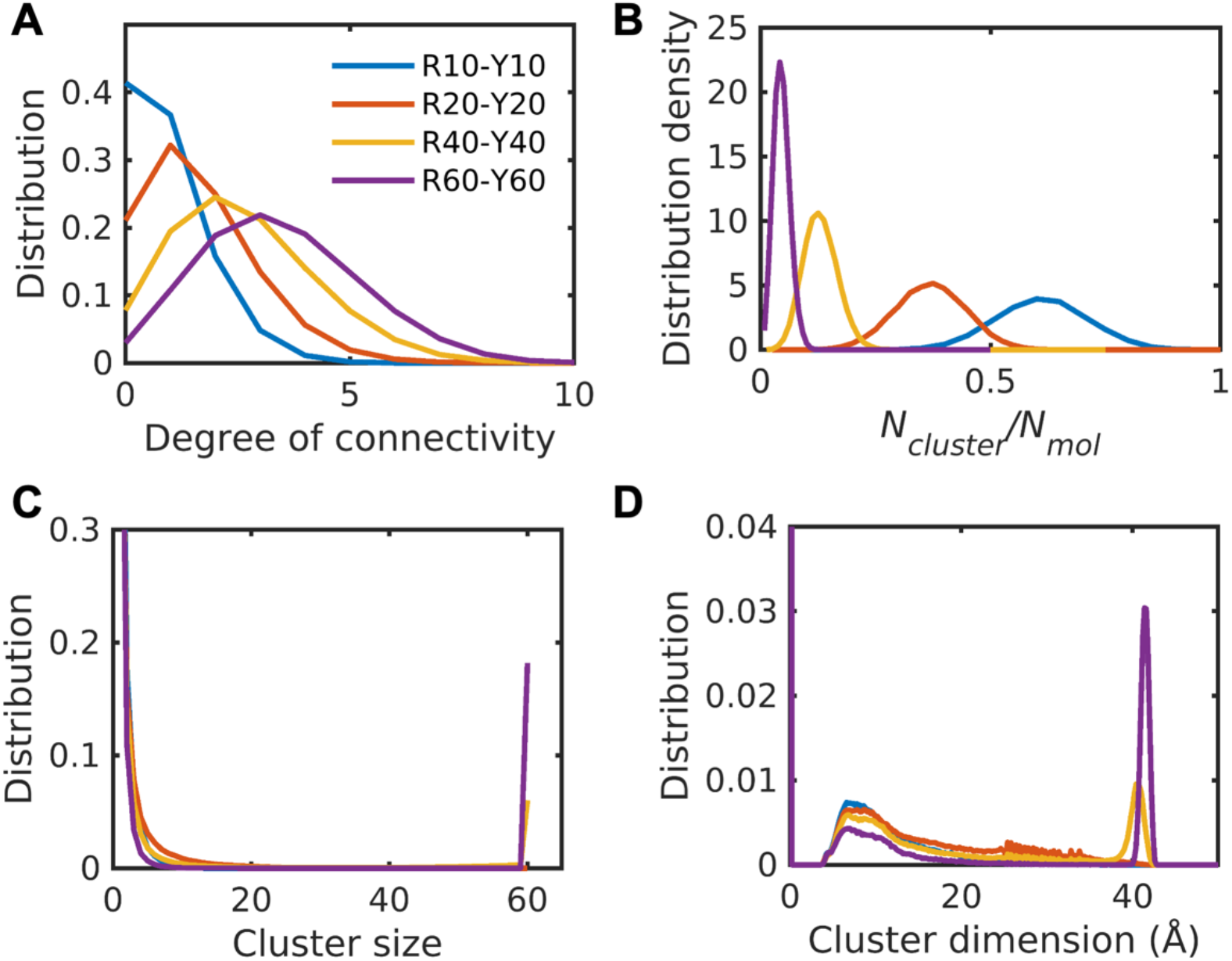
Dense phases modeled using capped amino acids form percolated networks. Distribution of degree of connectivity of each capped amino acid (A), distribution density of number of clusters per molecule (B), and distribution of cluster size (C) and cluster dimension (D) within the dense phase. The degree of connectivity, cluster size, and cluster dimension increase with increasing concentration, whereas the number of clusters per molecule decreases.

Next, we analyzed the dynamics of contacts that form and break within percolated networks. First, the timescales of amino acid contacts were quantified using the survival probability, defined as the conditional probability that a contact that was observed at time *t* persists at time *t*+*dt*. The results are shown in **Figure S9**. The half-life of a contact was determined as the time point where the survival probability equals 0.5. This defines a stochastic separatrix ^86^ and the choice of 0.5 was inspired by the *pfold* order parameter that has been used in characterizations of transition state ensembles in protein folding simulations ^87, 88^. The calculated half-lives for Y-Y, R-Y and R-R contacts are 0.48 ns, 0.27 ns and 0.07 ns, respectively. These results indicate that Y-Y interactions have longer lifetimes than R-Y interactions, and these are stronger than R-R interactions. The half-lives extracted from survival probabilities are consistent with the PMFs. These results confirm that the contact lifetimes are determined by specific amino acid interactions rather than stochastic contacts that might be of higher likelihoods at higher mass concentrations.

We also analyzed the temporal evolution of the network structure by analyzing how the average degree of connectivity and the largest cluster size vary as a function of time. Both the average degree of connectivity and the largest cluster size increase during the first 10 ns and then fluctuate around equilibrium values (**Figure S15**). These results demonstrate that beyond an equilibration / relaxation timescale of 10 ns, the network rearranges via an equilibrium flux of associations and dissociations.

## Discussion

In this work, we used all-atom simulations based on the AMOEBA polarizable forcefield to quantify differences between pairwise associations of aromatic and cationic sidechains in mimics of dilute versus dense phases of protein-based condensates. The design of the simulations leverages the fact that protein concentrations in dense phases place these systems in the semidilute regime ^89^ that are at ^28^ or above the overlap concentration ^9, 10, 28, 90^. These semidilute solutions are good or athermal solvents and the hence the number of residues within a concentration blob should be close to one. This ensures that our results are directly relevant to protein condensates.

For freely diffusing model compounds, the simulations show evidence of long-range ordering within dense phases that leads to the presence of nanoscale aromatic clusters and dispersed cationic compounds, ions, and water molecules. The hierarchies of interaction strengths are in accord with recent energy decompositions ^91^. In our simulations, which are based on the AMOEBA forcefield ^49^, the pairwise associations are enhanced in dense phases compared to dilute phases. In contrast, for simulations of capped amino acids, we find that backbone-mediated interactions weaken pairwise associations between sidechains, and this causes a loss of long-range ordering, and disruption of the nanoscale clusters of aromatic moieties. Fluidization is evident in the loss of long-range ordering. The increased valence afforded by the presence of peptide backbones, taken together with the weakening of pairwise sidechain associations, helps drive the formation of system-spanning networks. These networks are defined by the totality of directional, sidechain-sidechain, sidechain-backbone, and backbone-backbone interactions.

Our simulations paint a new picture regarding the driving forces for phase separation of PLCD-like systems: Strong pairwise associations between sidechains ^73^ are present in dilute phases, and these, as shown experimentally, are direct determinants of the free energy of transfer of polypeptides into dense phases ^25^. Strong pairwise interactions ^92^ are consistent with the designation of aromatic and Arg residues as stickers ^4, 25^. In fact, the intrinsic interaction energies are further enhanced in dense phase mimics based on model compounds, thus further corroborating the designation of these residues as stickers ^92, 93^. The favorable inter-residue two-body interactions outcompete chain-solvent interactions and the entropy of mixing, thereby driving phase separation^94^. However, within dense phases, the interaction patterns are different from the dilute phase. This is true for model compounds and for peptides. For model compounds, the lack of a backbone enables a range of rotations and translations of the compounds, thereby engendering long-range ordering and the formation of nanoscale clusters that coexist with regions that are devoid of these clusters. Backbones interfere with this clustering by introducing extra valence that weaken pairwise associations between sidechain moieties. Weakening of pairwise associations enables networking and drives percolation within dense phases. Indeed, recent studies ^60^ have shown evidence for condensates being percolated network fluids ^10, 27, 81, 95, 96^. Furthermore, the network structures directly explain the measured, sequence-specific viscoelastic moduli of different PLCD- based condensates ^27^.

Our studies lead to two key findings: First, the interactions within dilute and dense phases are quantifiably different and these differences are expected to be generators of interphase properties that include viscoelasticity ^27, 56^ and the generation of interphase electrochemical potentials ^12, 97^. Second, while dense phases of small molecules such as model compounds are likely to drive crystallization transitions ^98^, as evidenced by the nanoscale clustering and apparent phase separation within dense phases of model compounds, the peptide backbone intervenes with competing directional interactions such as hydrogen bonds to fluidize dense phases.

To investigate changes to solvent properties, we compared the induced dipole moments of water molecules and the model compounds mimicking the side chain of Tyr in both phases. The computation of induced dipole moments requires the use of a polarizable forcefield. We found that the dipole moments for water molecules in the dilute phase and dense phase are 2.71±0.23 Debye, and are identical across the phase. We obtain a similar result for the static, induced dipole moment of water molecules, which 2.65±0.52 Debye in both phases. These results suggest that the intrinsic physiochemical properties of the molecules remain unchanged between the dilute and dense phases. Instead, the differences between dense and dilute phases arise primarily from the effects of crowding in the dense phase. Notably, the extent of hydration of amino acids, quantified by analyzing water molecules around the amino acids, is diluted in the dense phase (**Figure S13**). Furthermore, the crowded environment does the dynamics of water molecules. Specifically, the self-diffusion coefficient of water in the dense phase is 1.30 × 10^-5^𝑐𝑚^2^/, compared to 2.10 × 10^-5^𝑐𝑚^2^/𝑠 in the dilute phase (**Figure S13**). These results are consistent with the findings of Mukherjee et al ^65^, who showed that hydration waters have longer residence times around proteins in the dense phase.

In both dilute and dense phases, there is a clear hierarchy of pairwise interactions, and there is a positive correlation between these hierarchies. This explains why models ^99^ based on transferrable ^99, 100^ or learned potentials ^9, 10^ that are pairwise in nature provide accurate descriptions of phase separation driven by homotypic and heterotypic interactions. Our dilute phase calculations are concordant with recent quantum mechanical energy decompositions ^91^. However, there are specific deviations vis-à-vis inferences made based on PMFs computed using non-polarizable forcefields ^18, 36^. These studies use a reaction coordinate that is the center-of-mass between the compounds of interest, whereas we used distances between functional groups of sidechains. Our choice is motivated by the formal definition of stickers, which implicates directional interactions between functional groups ^92, 94^. Converting PMFs from simulations that use non-polarizable forcefields and use centers-of-mass as the reaction coordinates, we derive, based on excluded volumes, the following interaction hierarchy in dilute phases: R-Y > Y-Y > R-F > F-F > K-Y > K-F. This contrasts with the hierarchies we derived from dilute phases of model compounds and capped amino acids, according to which: W′-W′ > R′-W′ > Y′-Y′ > F′-F′ > R′-Y′ > R′-F′ ≈ K′-Y′ > K′-F′. The key discrepancy pertains to the relative strengths of R-Y versus Y-Y and R-F versus F-F interactions. The calculations based on the AMOEBA forcefield, which includes electronic polarizability for water molecules, ions, and the peptides, suggest that the effects of polarization make π-π interactions stronger than the Arg-π interactions. The inferences from our calculations are concordant with solubility data ^101^, and inferences from experimental data, which show that π-π interactions are likely to be intrinsically stronger than cation-π-π interactions with the caveat that these effects are also context dependent ^4, 9, 10, 15, 16, 17, 25, 40, 102, 103^. It is noteworthy that the entropy-energy decompositions suggest, in accord with Grimme ^73^ as well as Martinez and Iverson ^74^, that π-π associations might be driven mainly by entropic considerations akin to what one observations for saturated, non-aromatic ring systems. They are also consistent with observations from THz spectroscopy ^57^.

Our simulations suggest that dense phases are fluidized and networked by the interplay of solvent-mediated sidechain-sidechain, backbone-sidechain, and backbone-backbone interactions that is absent in solutions of model compounds. These findings emphasize the importance of backbone-mediated interactions as drivers of weakened pairwise interactions, decreased long-range ordering that enables fluidization, and percolation within dense phases ^104^. Taken together, our results also highlight the need for using coarse-grained models with explicit representations of polypeptide backbones as recently introduced by Zhang et al., ^105^. This method adapted previous approaches for coarse-graining that preserve the backbone geometry and interactions ^106, 107^. Alternatively, one can use all-atom descriptions if computational resources are available ^31^. However, as a versatile, computationally tractable, and robust method, the approach of Zhang et al., ^105^ holds considerable promise.

We used a polarizable forcefield, AMOEBA. This is justified based on the accuracy of AMOEBA, which is now well-established, and the desire to capture at least first-order induced dipole contributions. We asked if as non-polarizable forcefield would yield equivalent results. For this, we calculated PMFs for amino acids in the dilute phase using CHARMM36m protein force fields ^108^. The PMF curves obtained from CHARMM36m (**Figure S10**) exhibit similar patterns to those from AMOEBA. These include the locations of the minima and the appearance of secondary minima around 8.5 Å. The excluded volumes calculated from the two forcefields show a strong positive correlation, with a Pearson correlation coefficient of 0.82. The CHARMM36m results confirmed several key observations: (1) Tyr is a stronger sticker than Phe; (2) Arg forms stronger cation-π interactions with Tyr and Phe when compared to Lys; (3) And Tyr-Tyr interactions are stronger than Arg-Tyr interactions. Note that CHARMM36m predicted relatively weaker Phe-Phe interactions when compared to Arg-Tyr. However, the overall results remain consistent with those obtained using the AMOEBA forcefield. In the dense phase mimics modeled using capped amino acids and the CHARMM36m forcefield, multivalent interactions between amino acids, including sidechain-sidechain, backbone-backbone, and sidechain-backbone interactions, drive the formation of the system-spanning network and weaken the pairwise interactions between sidechains. Overall, our findings suggest that that the choice of forcefields do not materially alter the major conclusions that emerge from our work.

In conclusion, we leveraged the fact that dense phases are semidilute solutions and used dense phases of peptides to understand the nature of interactions within condensates. In the blob picture, chain connectivity primarily imposes topological constraints on amino acids, and the dense phase can be approximated as concentrated solutions of peptide-sized motifs. Therefore, our results represent a first step toward uncovering differences in pairwise interactions, at atomic resolution, between dilute and dense phases. The formation of percolated networks within dense phases, are likely to be amplified when considering a semidilute solution of full-length chains.

However, the nature of the physical crosslinks that drive percolation are likely to be equivalent to what we have presented in our work.

## Methods

### Setup of molecular dynamics simulations using AMOEBA polarizable forcefield

All the simulations were performed using TINKER 9 package ^109^ employing the AMOEBA polarizable forcefield ^49^. The size of the central simulation cell, which was cubic, was ∼ 5 × 5 × 5 nm^/^ for both the dilute and dense phase simulations. The forcefield parameters for model compounds mimicking sidechains of Arg, Lys, Tyr and Phe were taken from our previous work ^110^. The forcefield parameters amoebabio18.prm in the TINKER package were used for capped amino acids. The isothermal-isobaric (NPT) ensemble was used. The temperature and pressure were maintained at 298 K and 1 bar, respectively. Other setup details about MD simulations were identical to our previous work ^110^. The model compounds or peptides were solvated in a cubic water box with periodic boundary conditions. Following energy minimization, molecular dynamics simulations were performed using the reversible reference system propagator algorithm integrator ^111^ with an inner time step of 0.25 ps and an outer time step of 2.0 fs in isothermal-isobaric ensemble (NPT) ensemble with the target temperature being between 273 and 400 K depending on the temperature of interest and the target pressure being 1 bar. The temperature and pressure were controlled using a stochastic velocity rescaling thermostat ^112^ and a Monte Carlo constant pressure algorithm ^113^, respectively. The particle mesh Ewald (PME) method ^114^ with PME-GRID being 36 grid units, an order 8 *B*-spline interpolation ^115^, with a real space cutoff of 7 Å was used to compute long-range corrections to electrostatic interactions. The cutoff for van der Waals interactions was set at 12 Å. This combination of a shorter cutoff for PME real space and longer cutoff for Buffered-14-7 potential has been verified ^116^ for AMOEBA free energy simulations ^117^. Snapshots were saved once every ten ps.

### Calculation of PMFs in the dilute phase using umbrella sampling

For the dilute phase simulations, a pair of model compounds or capped amino acids were placed in the simulation box. Umbrella sampling ^118^ was used to calculate the PMFs of interest. For the dense phase simulations, multiple pairs of model compounds or capped amino acids were placed in the simulation box. The number of pairs were estimated based on the volume fractions of dense phases gleaned from measured binodals of A1-LCD and designed variants thereof ^25^. Five independent simulations with different starting structures were performed for simulations that mimic dense phases. Each independent simulation was 100 ns long.

The interactions between pairs of model compounds or capped amino acids in the dilute phase were calculated using umbrella sampling. A harmonic potential with 𝑘 = 0.5 kcal/mol-Å^2^ was applied to restrain the distance between pairs of model compounds or capped amino acids. The geometric center of the key functional group in the model compound or the side chain of the capped amino acids was used to calculate distances along the reaction coordinates. For Arg and Lys, the key functional groups are the guanidinium and ammonium groups, respectively. For Tyr and Phe, the aromatic ring is the key functional group.

The centers of the windows for each umbrella are 3 Å, and 4 Å to 20 Å with an interval of 2 Å. In each window, multiple independent simulation trajectories with different starting structures are performed. The total simulation length for each window ranges from 240 ns to 260 ns. The weighted histogram analysis method (WHAM) ^119, 120, 121^ is used to obtain PMFs from umbrella sampling simulations. The distributions of pairwise distances, *P*(*r*) were first extracted from the simulations. These were then used to compute the PMFs of interest using: (𝑟) = 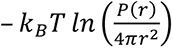. Here, *k_B_* is the Boltzmann constant, and *T* = 298 K is the simulation temperature.

### Estimation of error bars

For the error estimation of excluded volume in the dilute phase, we used bootstrapping with replacement. First, 160 ns of simulation data in each umbrella sampling window was chosen randomly to calculate the PMF for a specific pair of model compounds or capped amino acids. The PMF was further used to calculate the excluded volume. Next, the above process was repeated ten times, resulting in ten estimates of the excluded volume. The standard deviation of these ten estimates were the statistical uncertainties for the excluded volume.

For the error estimation of excluded volume in the dense phase, five values of the excluded volume can be calculated using each of the five independent simulation trajectories. Then, the standard deviation of these five estimated values was used to represent the error bars for excluded volume.

### Evaluations of convergence

To assess convergence, we performed a percolation analysis using one-fifth of the total simulation data. The results as shown in **Figure S14** match those obtained using the complete dataset in **Figure 10**, indicating that structural reorganization has reached convergence. Additionally, we plotted the average degree of connectivity and the largest cluster size as a function of time. As shown in **Figure S15**, both the average degree of connectivity and the largest cluster size increase during the first 10 ns and then fluctuate around their equilibrium values. These results demonstrate that amino acids continuously associated and dissociated, and the system achieves equilibrium after 10 ns.

## Author Contributions

Conceptualization: XZ and RVP; Methodology: XZ and RVP; Analysis: XZ and RVP; Funding acquisition: RVP; Project administration: RVP; Supervision: RVP; Writing: and Editing: XZ and RVP.

## Acknowledgments

This work was funded by the US Air Force Office of Scientific Research grant (FA9550-20-1- 0241 to RVP), the St. Jude Research Collaborative on the Biology and Biophysics of RNP Granules (to RVP), the US National Science Foundation (MCB-2227268 to RVP), and the Center for Biomolecular Condensates in the James McKelvey School of Engineering at Washington University in St. Louis.

## Additional Information

**Supplemental information** includes all supplemental figures, **Table S1 and Table S2**.

**Correspondence** and requests for materials should be addressed to Xiangze Zeng or Rohit Pappu.

