## Supplemental Figures and Tables for "Backbone-mediated weakening of pairwise interactions enables percolation in peptide-based mimics of protein condensates"

**for**

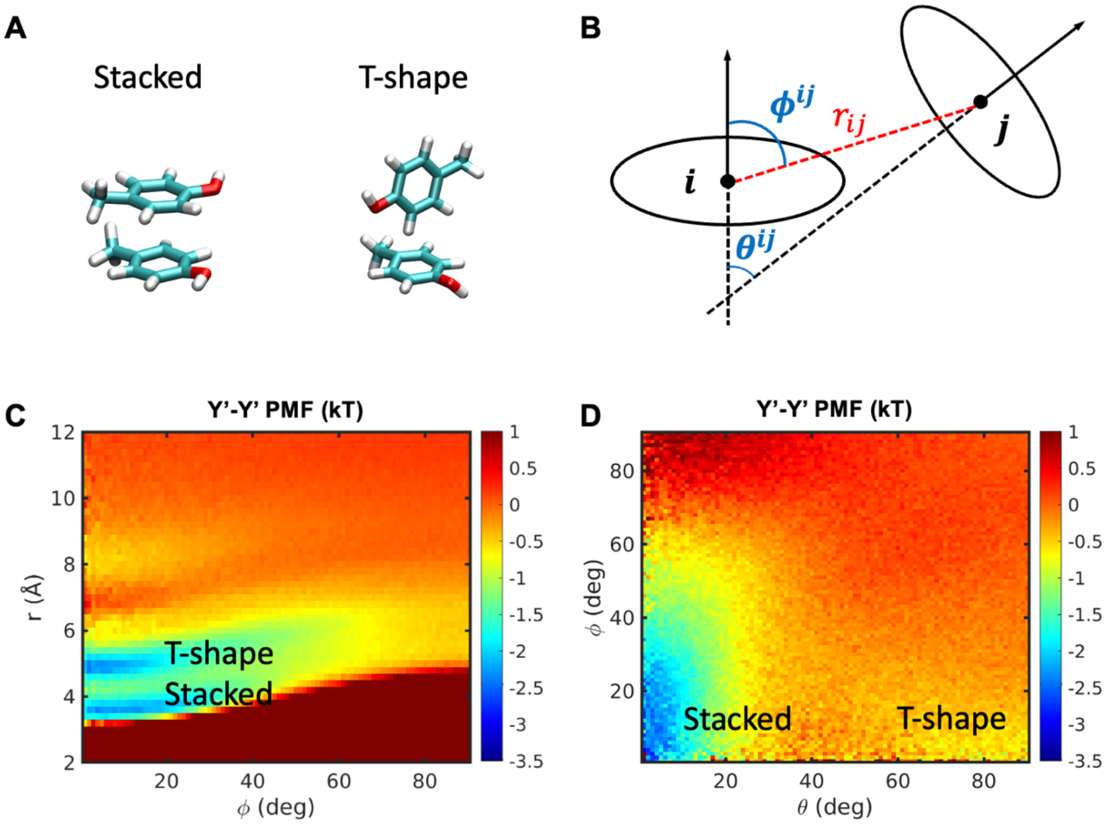


**Figure S1. Stacked and T-shape structures are formed by model compounds that mimic the side chain of Tyr.** (A) The stacked and T-shape structures. (B) Three parameters $r, \theta$ and $\phi$ were used to quantify the conformations of a pair of model compound mimics. $r_{ij}$ is the distance between geometry centers of key functional groups in the model compound $i$ and $j$. The key functional group are the aromatic ring for Tyr and Phe, the carbon atom in the guanidinium group for Arg, and the nitrogen atom in the ammonium group for Lys. For planar functional groups, we defined a plane by selecting three atoms to represent the orientation of the structure and $\theta$ is the angle between two normal vectors of the planar structures. Additionally, $\phi^{ij}$ is the angle between the vector $\vec{r}_{ij}$ and the normal vector of the planar structure i. These three order parameters allowed us to distinguish the stacked, parallel-displaced and T-shape conformations for π-π and cation-π interactions. The two-dimensional free energy profiles of a pair of Tyr model compound mimics projected onto (C) $r$ and $\phi$. (D) $\phi$ and $\theta$ show two metastable states representing the T-shape structure and stacked structure, respectively. The two-dimensional free energy profile is calculated from the dense phase simulations with 40 pairs of Arg and Tyr.

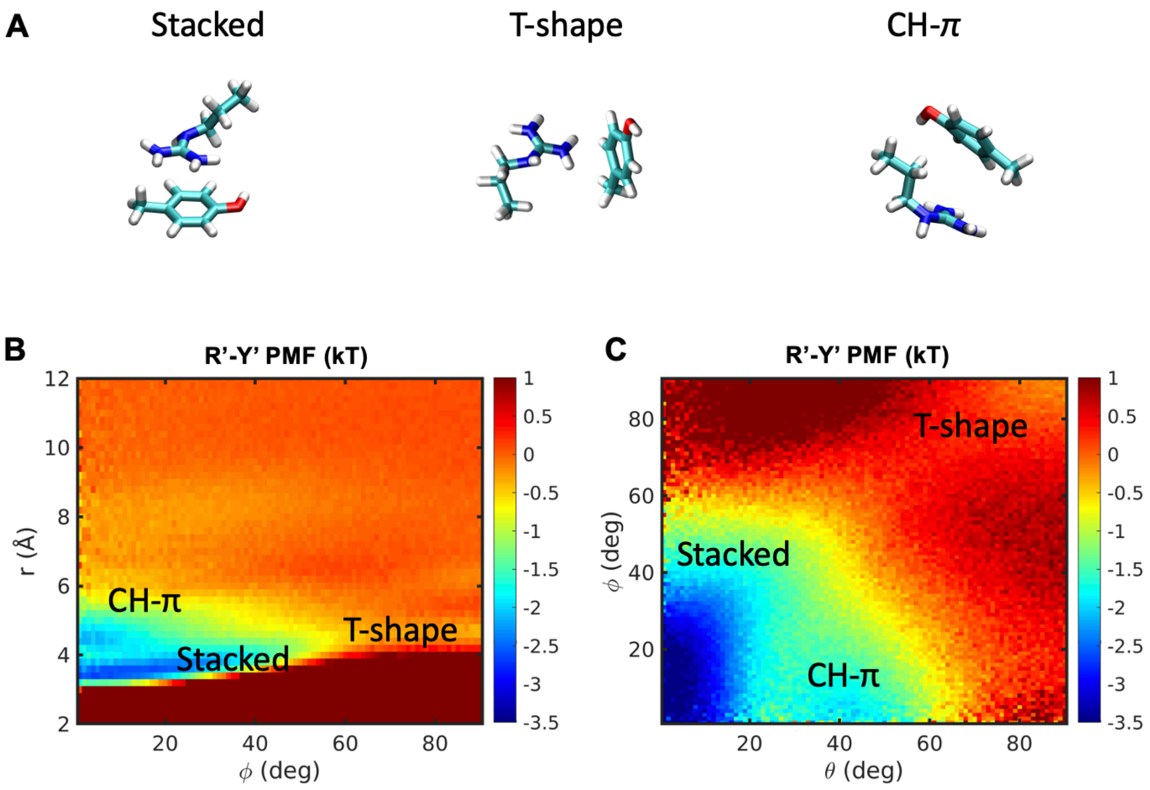


**Figure S2. Metastable structures are formed by model compounds that mimic the side chains of Arg and Tyr.** (A) The stacked structure, T-shape structure, and the structure showing the CH- $\pi$ interaction. The three order parameters defined in Figure S1B were used to quantify the conformations of a pair of Arg and Tyr model compound mimics. Here, the carbon atom in the guanidinium group of Arg model compound mimic was used as the origin of coordinates when calculating the three order parameters. The two-dimensional free energy profiles of a pair of Arg and Tyr model compound mimics projected onto (B) $r$ and $\phi$, and (C) $\phi$ and $\theta$ show metastable states representing the T-shape structure, stacked structure, and CH-$\pi$ interaction, respectively.

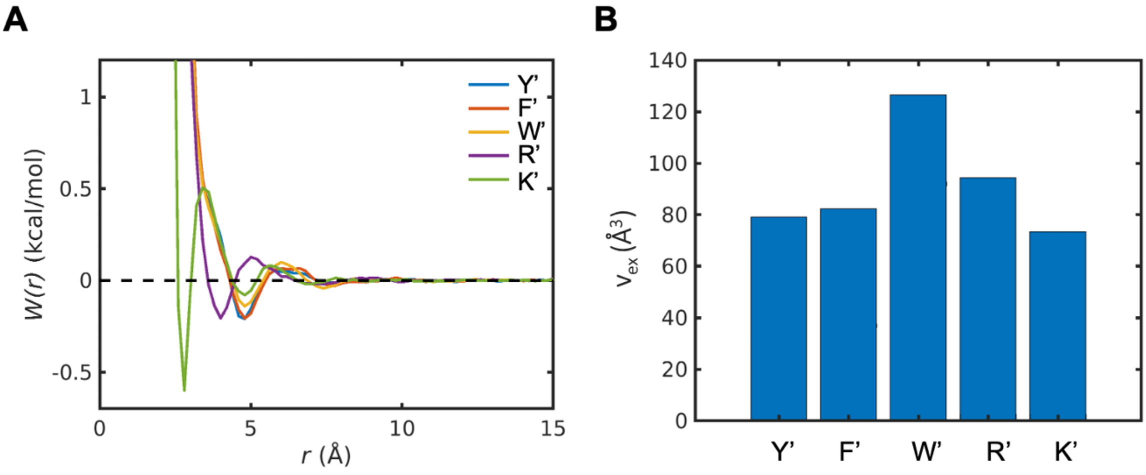


**Figure S3. Model compounds that mimic sidechains of amino acids have weak pairwise interactions with water molecules.** (A) PMFs, *W(r),* between the oxygen atoms in waters and key functional groups in the model compounds, which is calculated from the radial distribution of waters around these key functional groups. (B) Excluded volumes ($v_{ex}$) calculated from *W(r)*. The results are calculated from dense phase simulations.

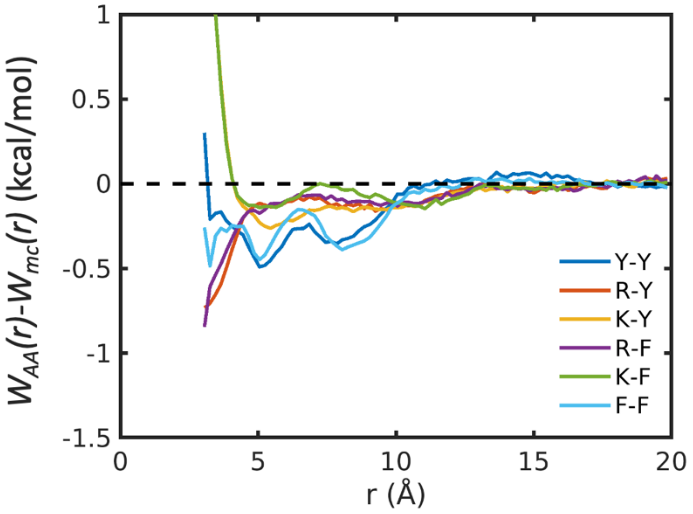


**Figure S4. The influence of backbone on interactions between side chains is side-chain-dependent.** The difference between the PMF for a pair of capped amino acids ($W_{AA}(r)$) and that for a corresponding pair of model compounds ($W_{mc}(r)$) is used to quantify backbone's influence.

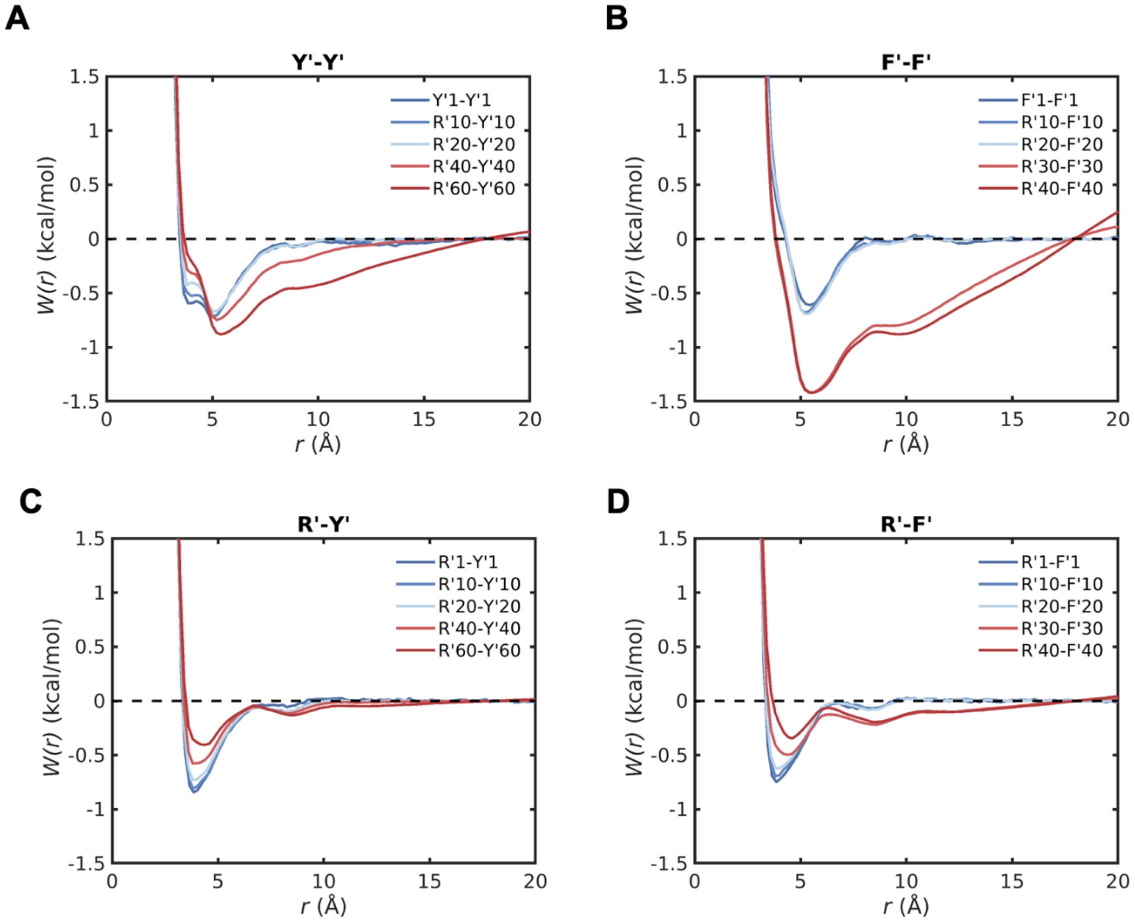


**Figure S5. Interactions among model compound mimics are weakened as increasing the concentration while maintaining a 1:1 ratio of Arg to aromatic moieties.** PMFs, *W(r)*, for (A) Y-Y, (B) F-F, (C) R-Y, and (D) R-F.

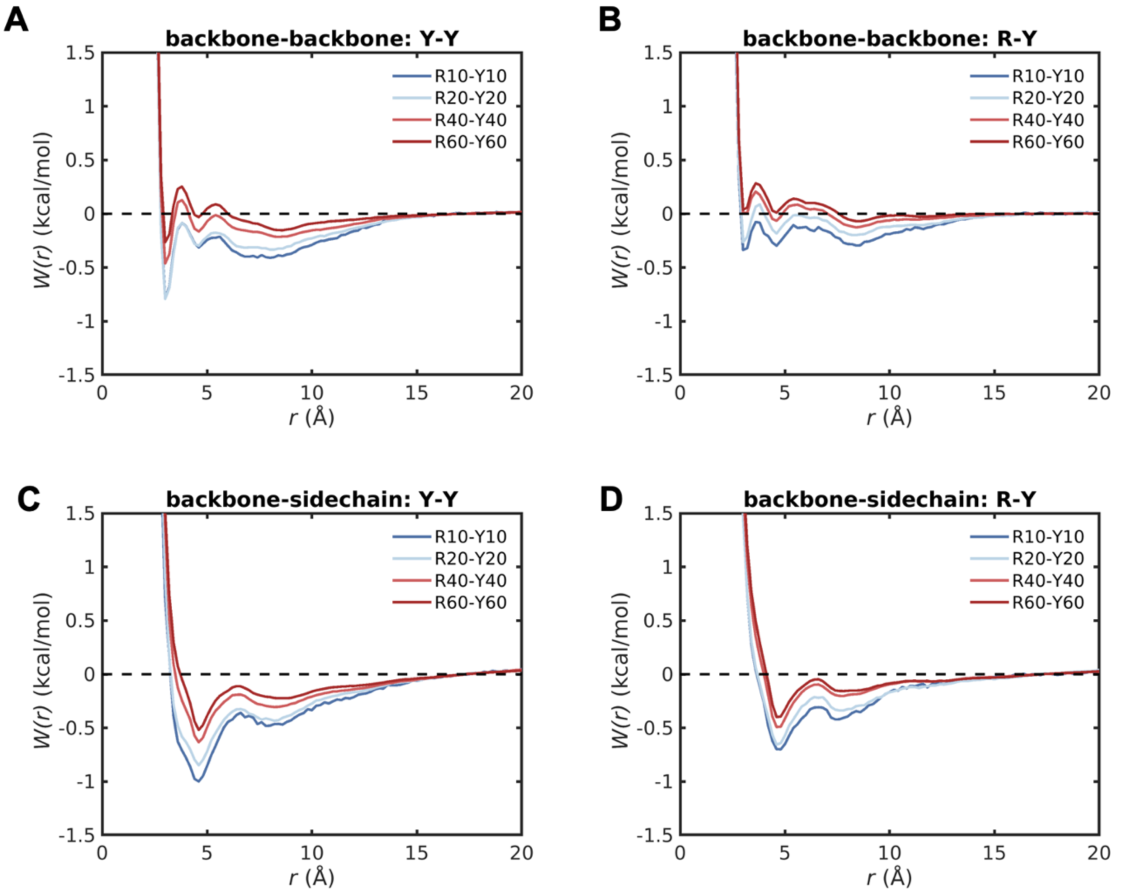


**Figure S6. Backbone-backbone and sidechain-sidechain associations among capped amino acids are weakened as increasing the concentration while maintaining a 1:1 ratio of Arg to aromatic moieties.** PMFs, *W(r)*, for (A) backbone-backbone interactions between Tyr, (B) backbone-backbone interactions between Arg and Tyr, (C) backbone-sidechain interactions between Tyr, and (D) backbone-sidechain interactions between Arg and Tyr. The backbone-backbone interactions are calculated using the backbone amide nitrogen atom and backbone carbonyl oxygen atom. The backbone-sidechain interactions were calculated using the backbone carbonyl oxygen atom and the geometric center of the carbon atoms in the aromatic ring of Tyr.

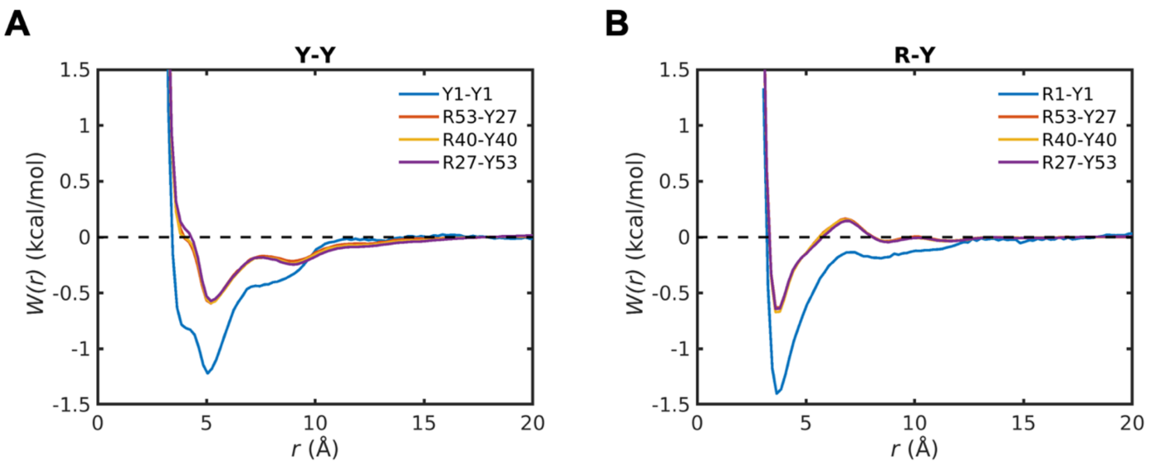


**Figure S7. The stoichiometry of Arg versus aromatic residues does not affect PMFs between capped amino acids in the dense phase.** PMFs, *W(r),* for (A) Y-Y, and (B) R-Y at different ratios of Arg to aromatic residues. Please note that Y1-Y1 and R1-Y1 indicate PMFs calculated from umbrella sampling in the dilute phase.

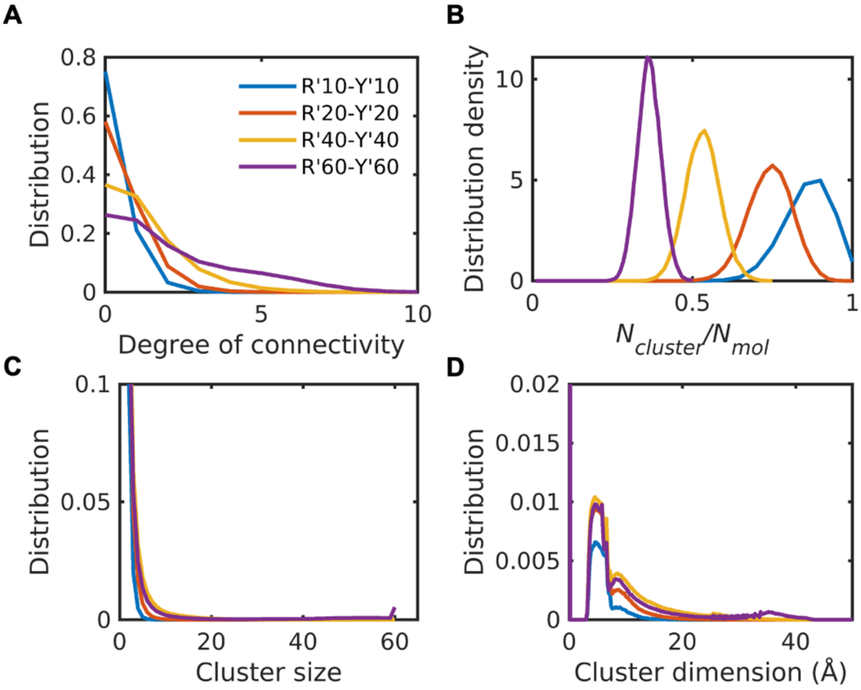


**Figure S8. Model compounds mimicking sidechains of amino acids in the dense phase do not form system-spanning network.** Distribution of degree of connectivity of each model compound (A), distribution density of number of clusters per molecule (B), and distribution of cluster size (C) and cluster dimension (D) within the dense phase. The degree of connectivity, cluster size and cluster dimension increase with increasing concentration, whereas the number of clusters per molecule decreases.

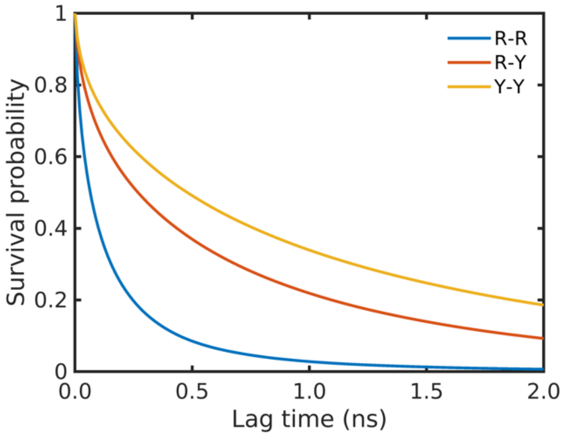


**Figure S9. R-R, R-Y and Y-Y contacts in the dense phase containing 40 pair of Arg and Tyr mixtures exhibit distinct half-lives quantified by survival probability.**

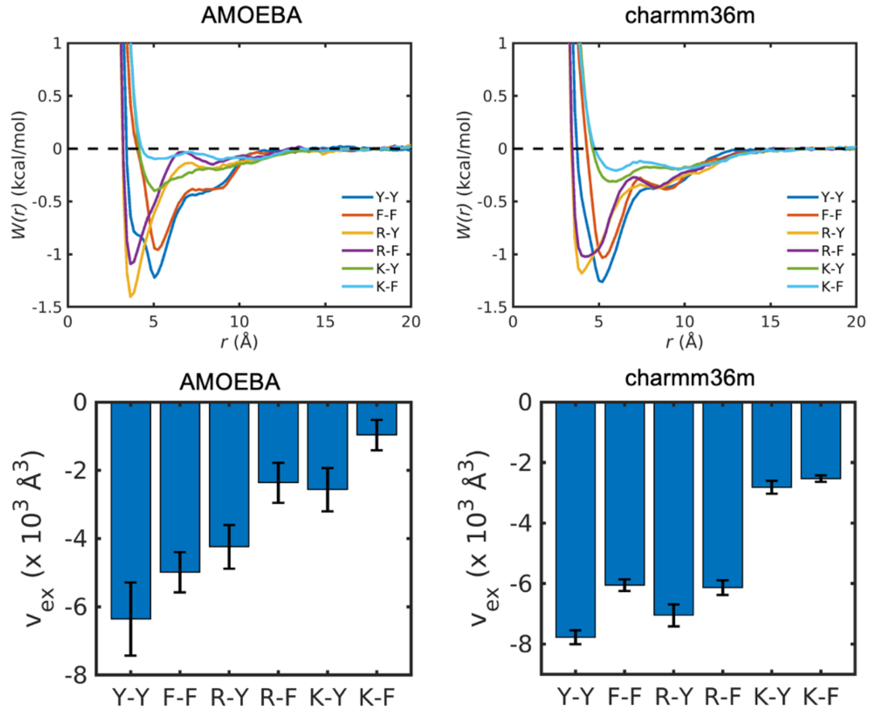


**Figure S10. PMFs (*W(r)*) for pairs of amino acids calculated from simulations using the polarizable AMOEBA and non-polarizable CHARMM36m force fields show very similar patterns.** The excluded volumes ($v_{ex}$) from these two force fields show a strong correlation with a correlation coefficient 0.82.

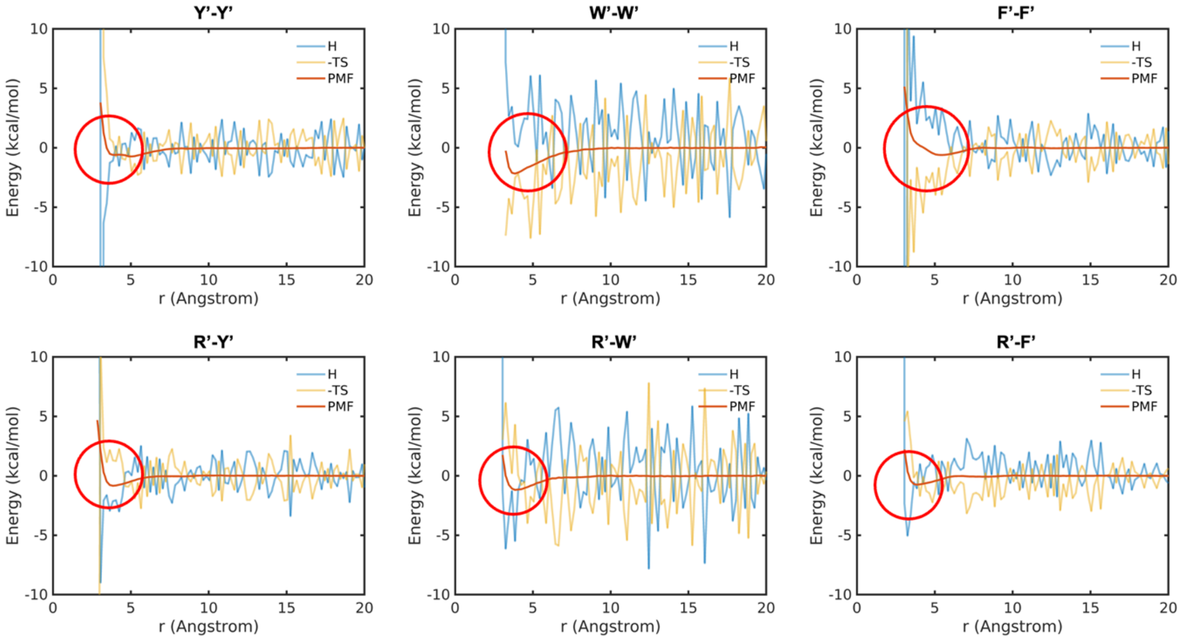


**Figure S11. Enthalpic and entropic contributions to PMFs for model compound pairs in the dilute phase.** For W'-W' and F'-F', entropy is favorable for the interactions. For other pairs, enthalpy drives the interactions at short distances (r < ~4Å), while the contributions at larger distances remain inconclusive due to significant fluctuations.

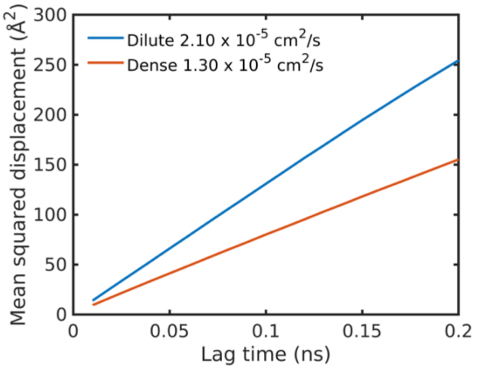


**Figure S12. The diffusion coefficient of water molecules in the dense phases is reduced due to interactions with amino acids, compared to that in the dilute phase.**

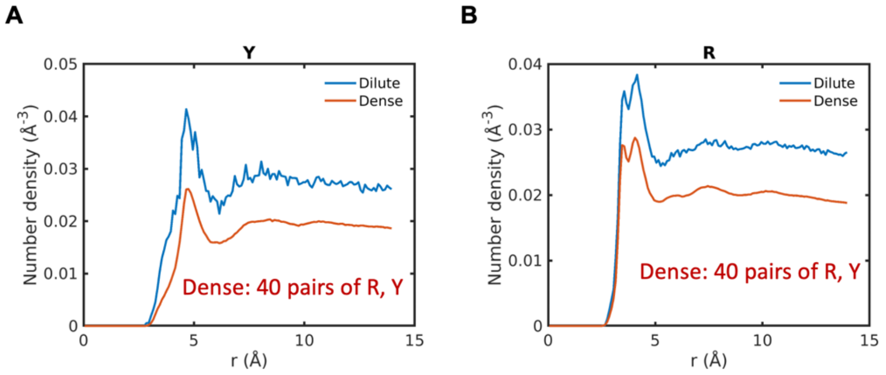


**Figure S13. Number density of water oxygen atoms around amino acids (A) Tyr and (B) Arg in the dense phase is lower than that in the dilute phase due to the crowding effect.**

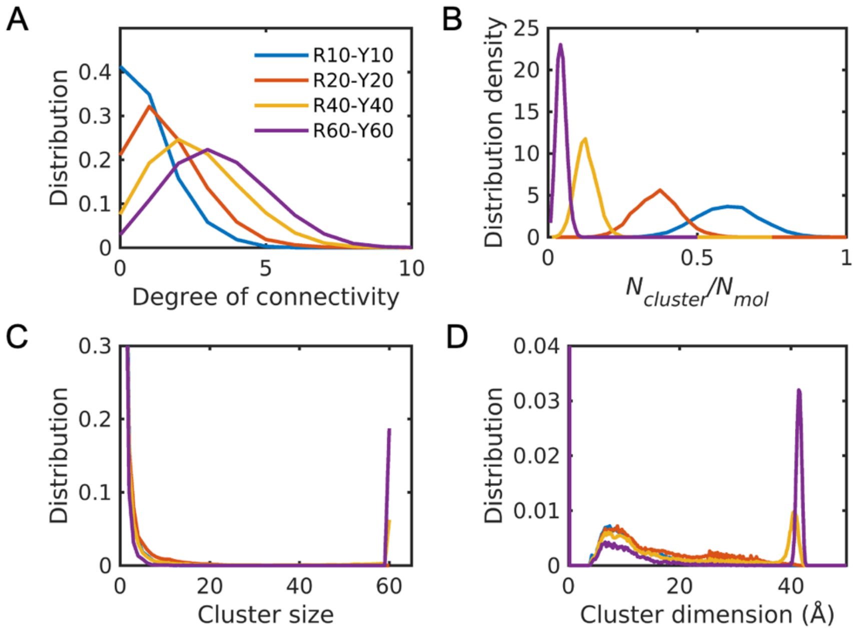


**Figure S14. Percolation analysis results using one fifths of the total simulation trajectories.** Distribution of degree of connectivity of each capped amino acid (A), distribution density of number of clusters per molecule (B), and distribution of cluster size (C) and cluster dimension (D) within the dense phase. The degree of connectivity, cluster size and cluster dimension increase with increasing concentration, whereas the number of clusters per molecule decreases. These results are nearly identical to those obtained from the full set of simulation trajectories as shown in Figure 10, demonstrating the convergence of the simulations.

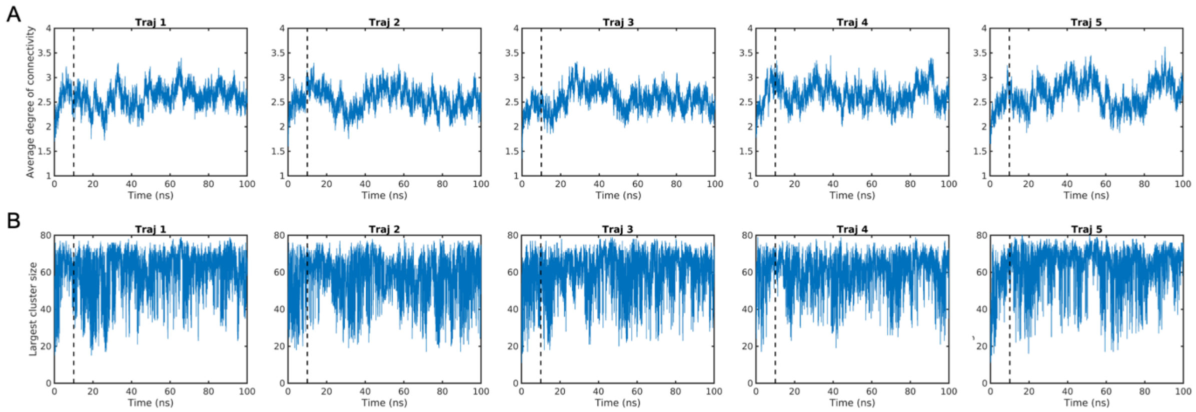


**Figure S15. Evolution of the average degree of connectivity of amino acids and the largest cluster size in the R40-Y40 mixtures.** The average degree of connectivity and the largest cluster size increase during the first 10 ns and then fluctuate around their equilibrium values.

**Table S1. Density of model compounds for different systems in this study.**

| | | | | | |
|---|---|---|---|---|---|
| **Model compounds mimicking amino acid sidechains** | **System** | Y’1-Y’1 | F’1-F’1 | W’1-W’1 | K’1-Y’1 |
| | ***Density (mg/mL)*** | *2.87* | *2.45* | *3.49* | *2.42* |
| | **System** | R’1-Y’1 | R’1-F’1 | R’1-W’1 | K’1-F’1 |
| | ***Density (mg/mL)*** | *2.79* | *2.58* | *3.10* | *2.21* |
| | **System** | R’40-Y’40 | R’40-F’40 | R’40-W’40 | / |
| | ***Density (mg/mL)*** | *111.75* | *103.25* | *123.99* | |
| | **System** | R’53-Y’27 | R’53-F’27 | / | / |
| | ***Density (mg/mL)*** | *110.71* | *104.98* | | |
| | **System** | R’27-Y’53 | R’27-F’53 | / | / |
| | ***Density (mg/mL)*** | *112.78* | *101.52* | | |
| **Capped amino acids** | **System** | Y1-Y1 | F1-F1 | W1-W1 | K1-Y1 |
| | ***Density (mg/mL)*** | *6.28* | *5.85* | *6.89* | *5.83* |
| | **System** | R1-Y1 | R1-F1 | R1-W1 | K1-F1 |
| | ***Density (mg/mL)*** | *6.20* | *5.99* | *6.50* | *5.61* |
| | **System** | R40-Y40 | R40-F40 | R40-W40 | / |
| | ***Density (mg/mL)*** | *247.92* | *239.42* | *260.15* | |
| | **System** | R53-Y27 | R53-F27 | K53-Y27 | K53-F27 |
| | ***Density (mg/mL)*** | *246.88* | *241.15* | *227.16* | *221.42* |
| | **System** | R27-Y53 | R27-F53 | K27-Y53 | K27-F53 |
| | ***Density (mg/mL)*** | *248.95* | *237.68* | *238.90* | *227.64* |

**Table S2. Distance cutoffs to determine the connectivity between molecules.** These cutoffs are positions of the first minimum of the radial distribution function for a specific pair of atoms or groups.

| | | |
|---|---|---|
| sidechain-sidechain | Aromatic ring to aromatic ring | 7.4 Å |
| | Aromatic ring to guanidinium group | 6.8 Å |
| | Guanidinium group to guanidinium group | 6.0 Å |
| sidechain-backbone | Aromatic ring to backbone carbonyl oxygen atoms | 6.6 Å |
| | Aromatic ring to terminal $CH_3$ group | 6.6 Å |
| | Guanidinium group to backbone carbonyl oxygen atoms | 5.0 Å |
| Backbone-backbone | Backbone amine nitrogen atoms to backbone carbonyl oxygen atoms | 3.8 Å |
